# A novel *cis*-element enabled bacterial uptake by plant cells

**DOI:** 10.1101/2022.03.28.486070

**Authors:** Chloé Cathebras, Xiaoyun Gong, Rosa Elena Andrade, Ksenia Vondenhoff, Jean Keller, Pierre-Marc Delaux, Makoto Hayashi, Maximilian Griesmann, Martin Parniske

## Abstract

The root nodule symbiosis (RNS) of plants with nitrogen-fixing bacteria is phylogenetically restricted to a single clade of flowering plants, which calls for yet unidentified trait acquisitions and genetic changes in the last common ancestor. Here we discovered – within the promoter of the transcription factor gene *Nodule Inception* (*NIN*) – a *cis*-regulatory element (*PACE*), exclusively present in members of this clade. *PACE* was essential for restoring infection threads (ITs) in *nin* mutants of the legume *Lotus japonicus*. *PACE* sequence variants from RNS-competent species appeared functionally equivalent. Evolutionary loss or mutation of *PACE* is associated with loss of this symbiosis. During early stages of nodule development, *PACE* dictates gene expression in a spatially restricted domain containing cortical cells carrying ITs. Consistent with its expression domain, *PACE*-driven *NIN* expression restored the formation of cortical ITs, also when engineered into the *NIN* promoter of tomato. Our data pinpoint *PACE* as a key evolutionary invention that connected *NIN* to a pre-existing symbiosis signal transduction cascade that governs the intracellular accommodation of arbuscular mycorrhiza fungi and is conserved throughout land plants. This connection enabled bacterial uptake into plant cells via intracellular support structures like ITs, a unique and unifying feature of this symbiosis.

## Introduction

Nitrogen is essential for plant growth and development ^1^. A wide phylogenetic variety of land plants ranging from mosses, gymnosperms to angiosperms have evolved symbioses with nitrogen-fixing bacteria that convert atmospheric nitrogen into ammonium ^2^. For example, the fern *Azolla* maintains colonies of nitrogen-fixing cyanobacteria in specialised apoplastic cavities, outside the plant cell wall enclosure ^3^. A major biological breakthrough was the evolution of the nitrogen-fixing root nodule symbiosis (RNS) characterised by the intracellular accommodation of bacteria in lateral organs (“nodules”) formed on roots ^4–6^. The occurrence of the RNS is restricted to a monophyletic clade, encompassing four angiosperm orders: the Fabales, Fagales, Rosales and Cucurbitales (FaFaCuRo) ^7^. Because of this phylogenetic restriction and scattered occurrence of RNS within the FaFaCuRo, Soltis and colleagues ^7^ postulated that the last common ancestor of the FaFaCuRo clade acquired a genetic change, a “predisposition”, which enabled members of this clade to subsequently evolve RNS multiple times independently ^7^. Intracellular accommodation of bacteria and root nodule development are two genetically separable and to this extend independent features of RNS ^8,9^. It is therefore genetically possible that they did evolve sequentially and not at the same time. The phylogenetic diversity of bacterial symbionts plus the variation of nodule anatomy and development across the RNS-competent FaFaCuRo species ^10,11^ together with the gap of 30 million years between the last common ancestor and the oldest fossil root nodules in this clade ^12^ further fuelled the hypothesis that nodule organogenesis evolved several times independently and was not a feature of the last common ancestor ^13,14^. The recent discovery of multiple losses of RNS within the FaFaCuRo clade ^15,16^ has initiated a discussion about whether this genetic change in the common ancestor was perhaps sufficient for the formation of RNS ^17^. Nonetheless, the precise nature of this key event in the evolution of nodulation has remained a mystery for more than two decades ^13^.

We asked which evolutionary acquisitions by the last common ancestor, in the form of novel traits and the underlying genetic causes, enabled the evolution of the RNS. From a phylogenetic perspective, such acquisitions should be: 1) exclusively present in the FaFaCuRo clade and absent outside of this clade and 2) conserved throughout the FaFaCuRo clade or at least maintained in RNS-competent (hereafter called “nodulating”) species. The uptake of bacteria into living plant cells is, with one exception (*Gunnera*), phylogenetically restricted to the FaFaCuRo clade ^4^. Uptake of bacteria requires the localised lysis of the plant cell wall, which threatens cell integrity because of the turgor pressure imposed by the protoplast ^6^. A systematic comparison of features associated with the RNS across the entire FaFaCuRo clade pinpoints a single unique and shared trait – the uptake of bacteria into living plant cells with intracellular physical support structures – that fulfils both abovementioned criteria to be acquired by the common ancestor ^6^. These structures come in a diversity of shapes (infection threads and infection pegs) and in at least two different cell types (epidermal and cortical), but are all characterised by the apposition of matrix material which is thought to maintain cell integrity during the localised lysis of the plant cell wall. While this matrix material is a common feature of all analysed successful bacteria uptake events in FaFaCuRo species, only one type, cortical infection threads (ITs), can be found in almost all nodulating species ^6^. Cortical IT formation is an evolutionary breakthrough because it allowed clonal selection of bacteria ^18^, specific control of nutrient exchange and increased nitrogen fixation efficiency ^19^. By contrast, in *Gunnera*, cell integrity is maintained by physical closure of a multicellular cavity by extracellular matrix material ^20^. This difference, together with the phylogenetic distance of *Gunnera* from the FaFaCuRo clade, suggests an independent origin of bacterial uptake in this genus ^6^. To search for gene gains specific for the FaFaCuRo clade, a genome-wide comparative phylogenomic analysis was performed, however, not a single gene following the aforementioned evolutionary pattern was identified ^15^.

Here, we tested the hypothesis that the “predisposition” event involved gain of novel *cis*-regulatory elements. Changes in gene regulation can be important drivers of functional and morphological evolution ^21,22^. Emergence or loss of even a single *cis*-regulatory element can lead to dramatic phenotypic consequences, e.g. novel organ formation ^21,22^. Phylogeny has dated the common ancestor of the FaFaCuRo clade to approximately 104 Mya ^23,24^. A long standing hypothesis states that the evolution of RNS involved co-opting genes from the arbuscular mycorrhiza (AM) symbiosis ^4,5^, which can be traced back to the earliest land plant fossils 410 Mya ^25,26^. This hypothesis is underpinned by similarities in intracellular accommodation structures ^6^ and the common requirement of both symbioses for a set of so-called “common symbiosis genes” ^5^ that are conserved across land plant species able to form AM, and encode symbiotic signal transduction and intracellular restructuring machineries ^27–31^.

## Results

### Discovery of *PACE*

The transcription factor-encoding *Nodule Inception* (*NIN*) gene ^32,33^ is positioned at the top of a RNS-specific transcriptional regulatory cascade and is indispensable for RNS ^32,34,35^. The promoter of *NIN* is a potential physical target for such a co-option event, because it defines the molecular interface between common symbiotic signal transduction and the specific transcriptional networks underlying RNS development ^35^. We therefore compared the *NIN* promoter sequences of 37 angiosperm species including 27 FaFaCuRo members and identified only one motif fulfilling the aforementioned criteria, which we called *Predisposition-Associated Cis-regulatory Element* (*PACE*) (**Fig. 1**; **Extended Data Fig. 1A – D** and **Supplementary Table 1**). The phylogenetic distribution of *PACE* was further investigated in an expanded search comprising 163 plant species in the promoter of *NIN* and the entire *NIN-like protein* (*NLP*) gene family, including *NLP1* from which *NIN* diverged at the base of the eudicots ^35^ (**Extended Data Fig. 1E** and **Supplementary Fig. 1**; **Supplementary Table 2**). *PACE* was found in all nodulating FaFaCuRo members and four non-nodulating species that have lost RNS but maintained *NIN* (**Extended Data Fig. 1E**; **Supplementary Table 3**). Importantly, *PACE* was absent from all the *NLP* promoters analysed (**Supplementary Fig. 1**). Thus, within the *NIN-like* gene family, the phylogenetic distribution of *PACE* is *NIN* and FaFaCuRo-clade specific and is consistent with a model in which *PACE* was acquired by the *NIN* promoter of the last common FaFaCuRo ancestor. Intriguingly, the 29 nucleotides-long *PACE* encompassed and extended beyond the previously identified binding site of the transcription factor Cyclops, encoded by a common symbiosis gene required for the development of both AM and RNS ^27,34^ (**Fig. 1** and **Extended Data Fig. 1**).

**Figure 1.**
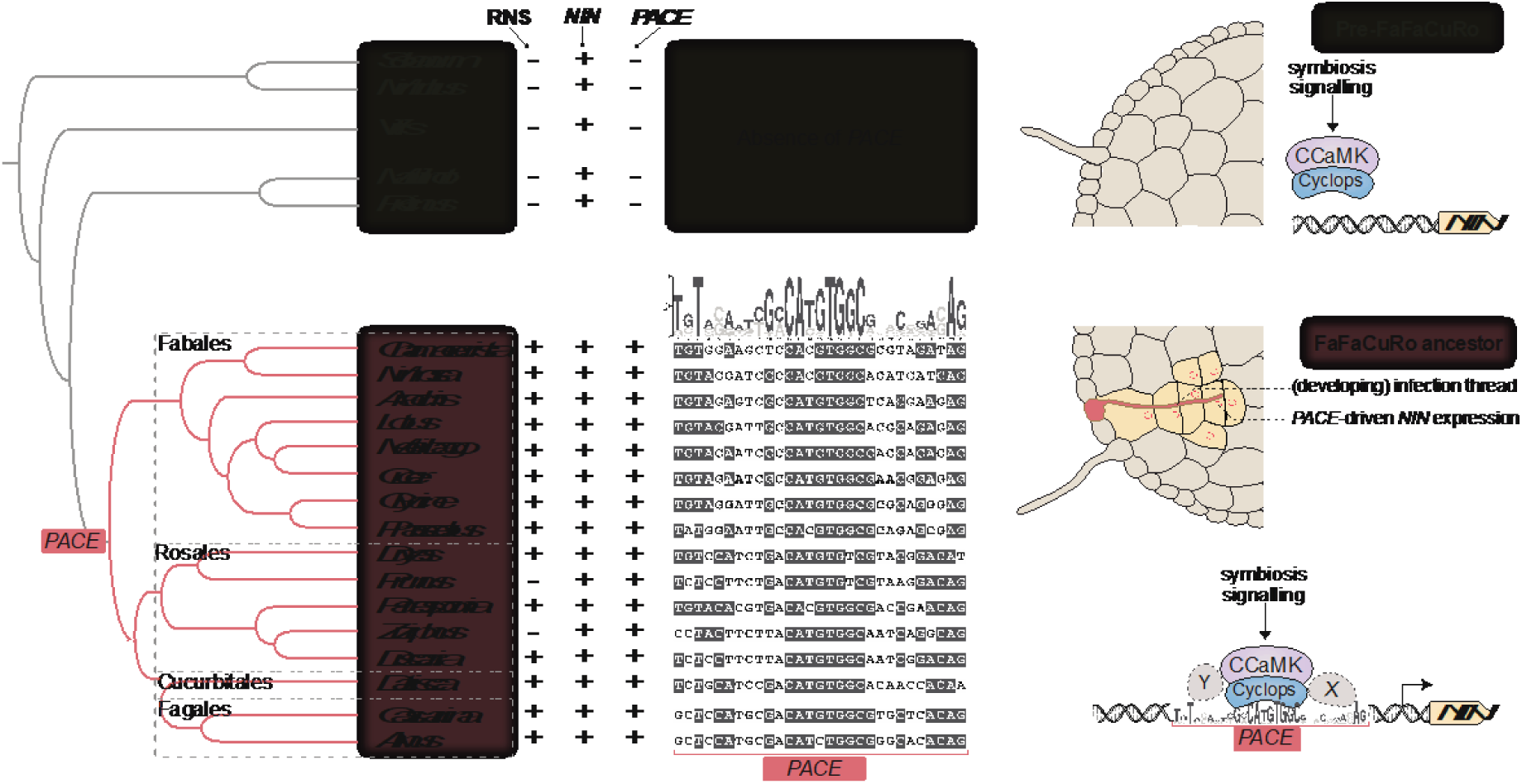
Acquisition of *PACE* was a key step in the evolution of RNS. **Left:** Schematic illustration of phylogenetic relationships between species inside (light red shade) and outside (light grey shade) the FaFaCuRo clade and presence (+) and absence (-) pattern of RNS, *NIN* and *PACE* (See **Extended Data Fig. 1 and Supplementary Fig. 1** for additional data support). **Middle:** *PACE* sequence alignment of the displayed species in which grey shadings indicate more than 50% sequence identity. On top of the alignment the *PACE* consensus sequence depicted as Position Weight Matrix calculated from the displayed RNS-competent species. Right: Graphical illustration of how *PACE* connected *NIN* to symbiotic transcriptional regulation by CCaMK/Cyclops, enabling IT development in the root cortex. This acquisition coincided with the predisposition event. X and Y: hypothetical proteins binding to sequences flanking the Cyclops binding site.

Given this clade-specific distribution of *PACE*, we searched for conserved motifs in the promoter sequences of two genes encoding transcriptional regulators, *ERF Required for Nodulation 1* (*ERN1*) ^36^ (**Supplementary Fig. 2**) and *Reduced Arbuscular Mycorrhiza 1* (*RAM1*) ^37^ (**Supplementary Fig. 3**) that are also known Cyclops targets. We identified motifs within the promoters of both, *ERN1* and *RAM1*, encompassing the previously identified Cyclops binding sites ^36,37^. In sharp contrast to *PACE*, their presence extended beyond the FaFaCuRo clade (**Supplementary Fig. 2** and **3**).

We tested the functional relevance of these distinct phylogenetic distribution patterns in transcriptional activation assays in *Nicotiana benthamiana* leaf cells. Transactivation by Cyclops was restricted to *NIN* promoters from FaFaCuRo species, but extended to non-FaFaCuRo species for *RAM1* promoters (**Extended Data Fig. 2** and **Supplementary Fig. 4**). Importantly, *PACE* was necessary and sufficient for the activation of the *NIN* promoter by Cyclops (**Extended Data Fig. 3**). Together with the exclusive occurrence of *PACE* in the *NIN* promoter of the FaFaCuRo clade, these results are in line with the hypothesis that the mechanistic link between Cyclops and the *NIN* promoter was established in the last common ancestor of this clade (**Fig. 1**).

### *PACE* drives the expression of *NIN* during IT development in the cortex

*NIN* is indispensable for IT development ^32,33^ and its precise spatiotemporal expression is essential for this process ^33,38–40^. Because *cis*-regulatory elements are master determinants of gene expression patterns ^41^, we investigated the impact of *PACE* on the expression of *NIN* in physical relation to the bacterial uptake and accommodation stages during nodule development. We used the model legume *Lotus japonicus* in combination with its compatible nitrogen-fixing bacterium *Mesorhizobium loti* as experimental system. The process by which *L. japonicus* promotes the intracellular colonisation by and accommodation of *M. loti* can be subdivided into successive stages: (1) entrapment of bacteria in a pocket formed by a curled root hair ^42^, (2) uptake of bacteria into a developing IT within that root hair ^42^, (3) IT progression into and through the outer cortical cell layers ^43^, (4) IT branching and extension within the nodule primordium ^44^ (5) release of bacteria from ITs into plant membrane-enclosed organelle-like structures called symbiosomes ^44^ leading to (6) mature nodules characterised by infected cells densely packed with symbiosomes and the pink colour of leghemoglobin ^45^.

To determine the *PACE-*mediated spatiotemporal expression domain, we introduced a *GUS* reporter gene driven by *PACE* fused to a region comprising the *NIN* minimal promoter and the 5’UTR ^34^ (*PACE:NINmin_pro_:GUS*) into *L. japonicus* wild-type roots. Roots were subsequently inoculated with *M. loti* MAFF 303099 expressing *Ds*Red (*M. loti Ds*Red) facilitating detection of the bacteria through their fluorescence signal in root hairs and nodules. The *NIN* minimal promoter did not mediate reporter gene expression at any stage of bacterial infection (**Extended Data Fig. 4E**). Intriguingly, the earliest detectable GUS activity mediated by *PACE:NINmin_pro_:GUS* was clearly restricted to a zone in the nodule primordia (panel I - II in **Extended Data Fig. 4D**) that roughly correlated with the site of bacterial infection (indicated by a local accumulation of *Ds*Red signal) and later expanded to the entire central tissue of the nodule (panel III in **Extended Data Fig. 4D**). *PACE*-driven reporter expression was neither detected in root hairs harbouring ITs (**Extended Data Fig. 4G**) nor in nodules in which cells from the central tissue were filled with symbiosomes (panel IV in **Extended Data Fig. 4D**). Importantly, *PACE-*mediated expression was distinct from that mediated by the *LjNIN* 3 kb promoter (*NIN_pr_*_o_) or the *NIN_pr_*_o_ with *PACE* mutated or deleted (*NIN_pro_::mPACE* and *NIN_pro_::ΔPACE*, respectively) that conferred reporter expression across the central tissue of the nodule (panels II - IV in **Extended Data Fig. 4A – C**). We concluded based on these observations that the *PACE-*mediated expression domain is temporally and spatially restricted and possibly accompanies the development of bacterial accommodation structures in the nodule.

To further resolve this relationship between *PACE* driven gene expression and bacterial accommodation at the cellular level, we compared – simultaneously in the same tissue – the progression of bacterial infection with the expression pattern mediated by *PACE* fused to the *NIN* minimal promoter (*PACE:NINmin_pro_*) and by a *NIN* promoter with mutated *PACE* (*NIN_pro_::mPACE*). A red and a yellow fluorescent protein (mCherry and YFP, respectively) targeted to the nucleus by fusion to a nuclear localization signal (NLS) were used as reporters. The resulting promoter:reporter fusions (*PACE:NINmin_pro_:NLS-mCherry* and *NIN_pro_::mPACE:NLS-YFP*) were placed in tandem on the same T-DNA allowing a nucleus-by-nucleus comparison of their relative expression. This T-DNA construct was introduced into *L. japonicus* wild-type roots that were subsequently inoculated with *M. loti* R7A expressing the cyan fluorescent protein (CFP; **Fig. 2**) or with *M. loti* MAFF 303099 expressing the green fluorescent protein (GFP; **Supplementary Fig. 5**) to facilitate detection.

**Figure 2.**
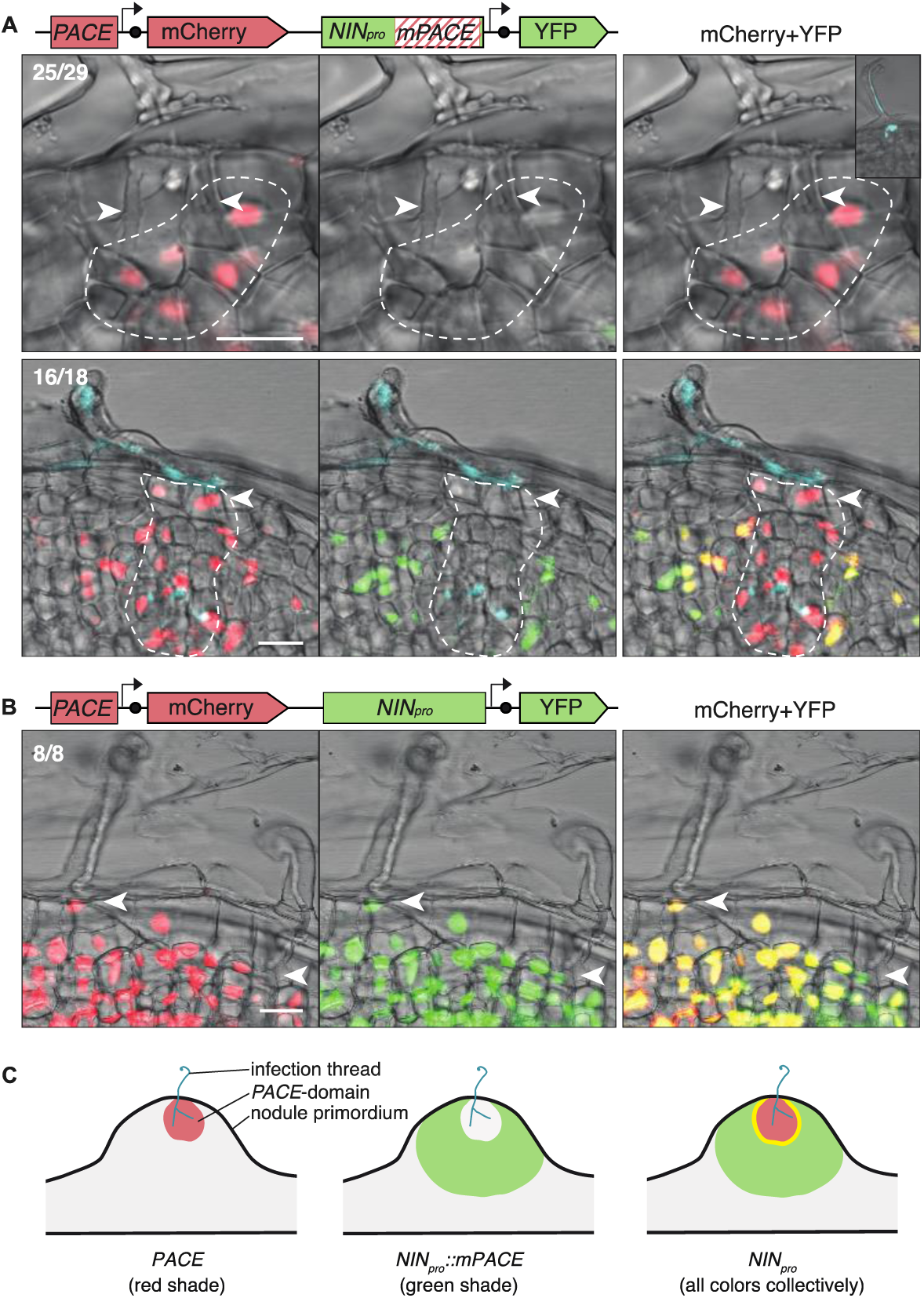
*PACE* drives the expression of *NIN* during IT development in the cortex. Sections of representative *L. japonicus* nodule primordia formed upon inoculation with *M. loti* R7A expressing CFP (blue) imaged by confocal laser-scanning microscopy. Comparison of the expression domains determined by (**A**) *PACE* (*PACE:NINmin_pro_:NLS-mCherry*; red) and a *NIN* promoter carrying a mutated *PACE* (*NIN_pro_::mPACE:NLS-YFP*; green); or (**B**) *PACE* (red) and the intact *NIN* promoter (*NIN_pro_:NLS-YFP*; green). Dashed lines demarcate a group of cortical cells in the *PACE* core territory. Arrowheads indicate ITs. Numbers: nodule primordia showing the presented expression pattern / total number of nodule primordia sectioned and inspected. Data are from four independent experiments (see **Supplementary Fig. 5** for the first stages of bacterial invasion (stage 2 to 3)). Bars, 20 µm. (**C**) graphical interpretation of expression patterns presented in (**A and B**). Yellow: overlapping region.

During the first stages of bacterial invasion (stage 2 to 3), *PACE*-mediated *mCherry* was expressed specifically in cortical cells carrying ITs and in directly adjacent cells (**Supplementary Fig. 5A**). By contrast, the *NIN_pro_::mPACE*-driven YFP signal was not detected in those cells (**Supplementary Fig. 5B**). In sections of developing nodules, in which infection had progressed to stage 3 or 4, *PACE*-mediated *mCherry* was expressed specifically in a – hereafter called “infection thread (IT) zone” – comprising cortical cells and primordium cells that carried ITs and in some, but not all, directly adjacent cells (25 out of 29 nodules inspected; **Fig. 2A**; ^46^). Intriguingly, the expression domains marked by mCherry and YFP fluorescence were distinct from each other: while the *PACE*-driven mCherry signal was consistently marking the IT zone, the *NIN_pro_::mPACE*-driven YFP signal was observed in primordium cells surrounding this zone (16 out of 18 nodules inspected; **Fig. 2A, 2C**). The thin (approx. 1-2 cells thick) border between the two domains was characterised by nuclei emitting both YFP and mCherry signals (**Fig. 2A**). In so-marked cells, ITs were typically not detected. The expression pattern mediated by the *NIN* promoter (containing *PACE*) was congruent with the sum of both promoter fragments (8 out of 8 nodules inspected; **Fig. 2B, 2C**).

Based on these clearly distinct and complementary reporter expression domains governed by *PACE* versus the remaining promoter, we concluded that 1) *PACE* directs *NIN* expression to a specific IT zone and that 2) the *NIN* promoter comprises *cis*-regulatory elements that drive expression outside the *PACE* territory i.e. in root hairs (together with *PACE*), non-infected cortical and primordium cells and nodule cells filled with symbiosomes. These additional *cis*-regulatory elements might be addressed by other transcription factors that have been reported to bind to this promoter ^47–49^. These transcription factors might be counteracted by, for example, repression in the IT zone.

### Mutational dissection of *PACE* reveals a quantitative impact of sequences flanking the *CYC-box* on IT development

To test the relevance and specific role of *PACE* in nodule and IT development, we performed complementation experiments using plants homozygous for the *nin-2* or *nin-15* mutant alleles ^32^. The *nin-2* mutant allele harbours a frameshift mutation of the *NIN* gene, leading to a *NIN* loss-of-function phenotype, i.e. absence of both IT formation and nodule organogenesis ^32^ while the *nin-15* mutant allele carries a *Lotus Retrotransposon 1* insertion within the *NIN* promoter 143 bp 3’ of *PACE* (**Extended Data Fig. 5**). We examined the restoration of bacterial infection 21 days after inoculation with *M. loti Ds*Red by quantifying the number of root hairs harbouring ITs and the number of infected nodules (**Fig. 3** and **Supplementary Table 4**).

**Figure 3.**
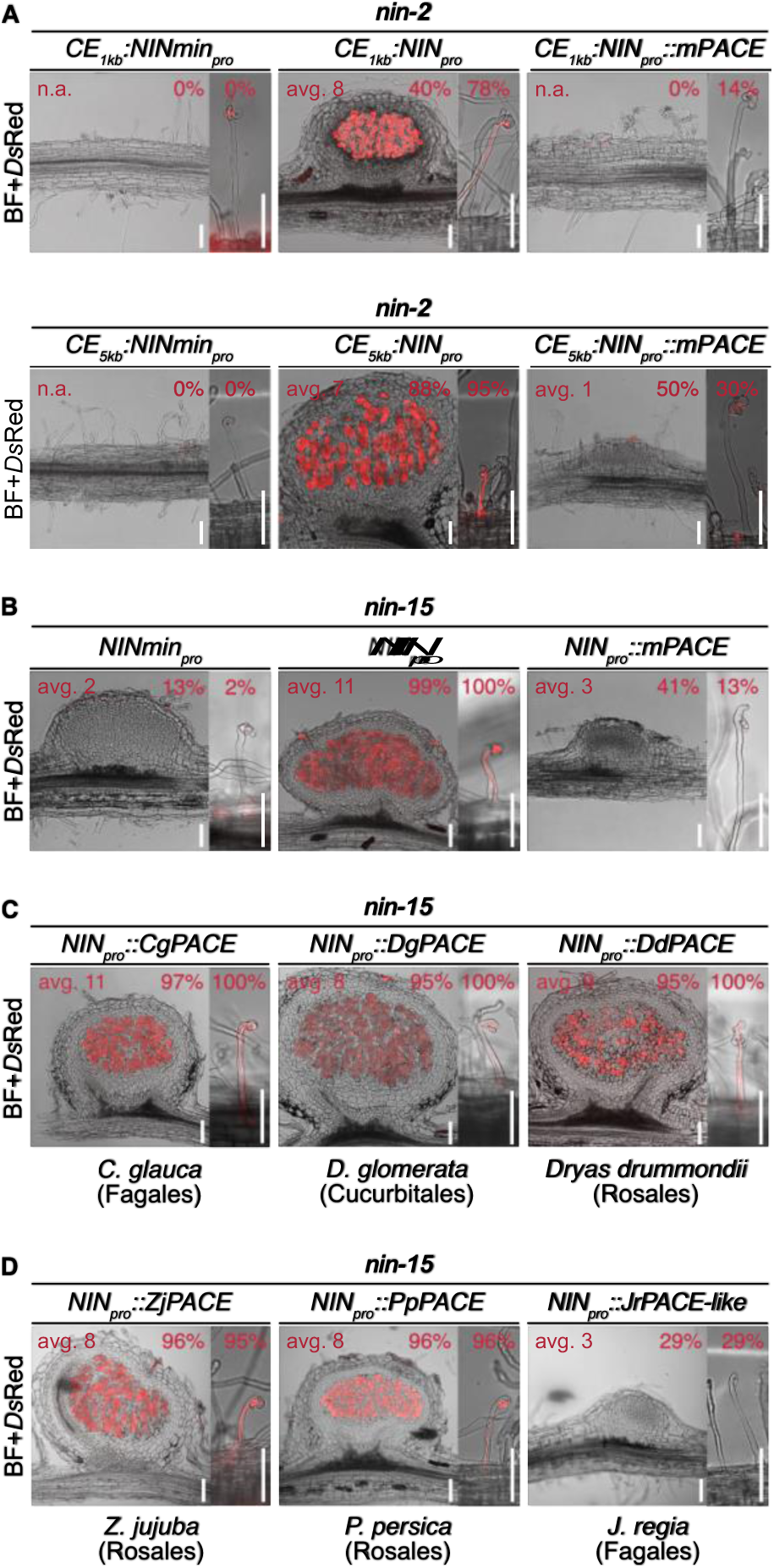
*PACE* is necessary for bacterial infection and functionally conserved across the FaFaCuRo clade. Microscopy images of representative nodule sections or root hairs harbouring an IT or an infection pocket from (**A**) *nin-2* or (**B - D**) *nin-15* roots transformed with the *LjNIN* gene driven by: (**A - B**) indicated promoters, and in (**C - D**) the *L. japonicus NIN* promoter in which *LjPACE* was replaced by *PACE* from (**C**) nodulating and (**D**) non-nodulating FaFaCuRo species or with a *PACE-like* sequence identified in the *JrNLP1b* promoter. %: percentage of transgenic root systems carrying infected nodules or root hair ITs. Avg.: average number of infected nodules on plants carrying infected nodules. n.a.: not applicable. At least five nodules from independent transgenic root systems were sectioned per construct. The percentage of root hair ITs among total infection events per root pieces and the number of infected nodules per transgenic root systems are displayed in **Extended Data Fig. 6 – 9**. BF, brightfield. Bars, 100 µm.

Nodule development in the legume *Medicago truncatula* is dependent on *NIN* expression mediated by a regulatory region containing several putative cytokinin responsive elements (*CE*) ^40^. In *L. japonicus*, a similar *CE* region is positioned 45 kb upstream of the *NIN* transcriptional start site ^40^. To enable transgenic complementation experiments, we synthetically fused a 1 kb or 5 kb region encompassing this distant *CE* to the 5’ end of a 3 kb *NIN* promoter. The *NIN* gene driven by these promoters (*CE_1kb_:NIN_pro_:NIN* and *CE_5kb_:NIN_pro_:NIN*) restored the formation of root hair ITs on 78% and 95% and infected nodules on 40% and 88% of *nin-2* transgenic root systems, respectively (**Fig. 3A**; **Extended Data Fig. 6 – 8** and **Supplementary Fig. 6 – 7**). Importantly, this complementation success relied on the presence of *PACE*; *nin-2* roots transformed with the same fusion design but carrying a mutation of *PACE* (*CE_1kb_:NIN_pro_::mPACE:NIN* and *CE_5kb_:NIN_pro_::mPACE:NIN*) did not restore root hair ITs but nodule formation was not impaired when using the cytokinin element-containing region of 5 kb (*CE_5kb_:NIN_pro_::mPACE:NIN*). We concluded that *PACE* is indispensable for bacterial infection but not for nodule development.

The 29 bp long *PACE* sequence revealed by MEME encompasses and extends beyond the previously identified Cyclops binding site (*CYC-box* ^34^, “box”, **Extended Data Fig. 1**). Its degree of conservation may be interpreted as a trace of an ancestral *PACE* version present in the last common ancestor of the FaFaCuRo clade. Within *PACE* the *CYC-box* is surrounded by less conserved flanking sequences. To dissect the specific contributions of the *CYC-box* and *PACE* sequences flanking the *CYC-box* (“flanking”) to *PACE* function, we mutated the box and the flanking sequences independently (*CE:NIN_pro_::mbox:NIN* and *CE:NIN_pro_::mflanking:NIN*, respectively). Mutation of the *CYC-box* abolished root hair ITs. Interestingly, mutation of the flanking sequences led to a 50% reduction of the number of transgenic root systems carrying infected nodules, while the formation of root hair ITs was not impaired (**Extended Data Fig. 6 – 8** and **Supplementary Fig. 6 – 7**). This mutational dissection revealed two separable functions of *PACE*: while the *PACE*-Cyclops connection is essential for IT development, the flanking sequences significantly promote bacterial infection during nodule development and possibly act as binding sites for additional, yet undefined, transcription factors (conceptually labelled X and Y in **Fig. 1**). Our data suggest that *PACE* comprises synergistic binding sites for both Cyclops and cooperating transcription factors. We conclude that the high level of conservation of the *CYC-box* is a consequence of the indispensable nature of this *cis*-element for the progression of the IT through the cortex. The higher level of diversification of sequences flanking the *CYC-box* might be a consequence of changes in transcription factors occupancy over evolutionary time scales. Considering this scenario, it is possible that such flanking sequence occupying transcription factors are not conserved throughout the entire FaFaCuRo clade.

*PACE*-mediated *NIN* expression defined an infection zone in the nodule cortex (**Fig. 2**). To genetically separate the initiation of nodule development from IT formation and thereby enable a focussed analysis of the role of *PACE* in cortical IT formation, we utilised the *nin-15* mutant, which is impaired in IT formation but retains the capacity to form nodules. Most of these nodules were uninfected (92% and 86% plants carrying no root hair ITs and no infected nodules, respectively) and cortical cells filled with symbiosomes were never observed (**Extended Data Fig. 5**). This mutant therefore provided an ideal background to study the role of *PACE* in cortical IT formation, circumventing the negative epistatic effect of the inability of *nin* loss-of-function mutants to initiate cell divisions ^32,38–40,50^ (**Fig. 3B – D**; **Fig. 4**; **Extended Data Fig. 9 – 10** and **Supplementary Table 4**).

**Figure 4.**
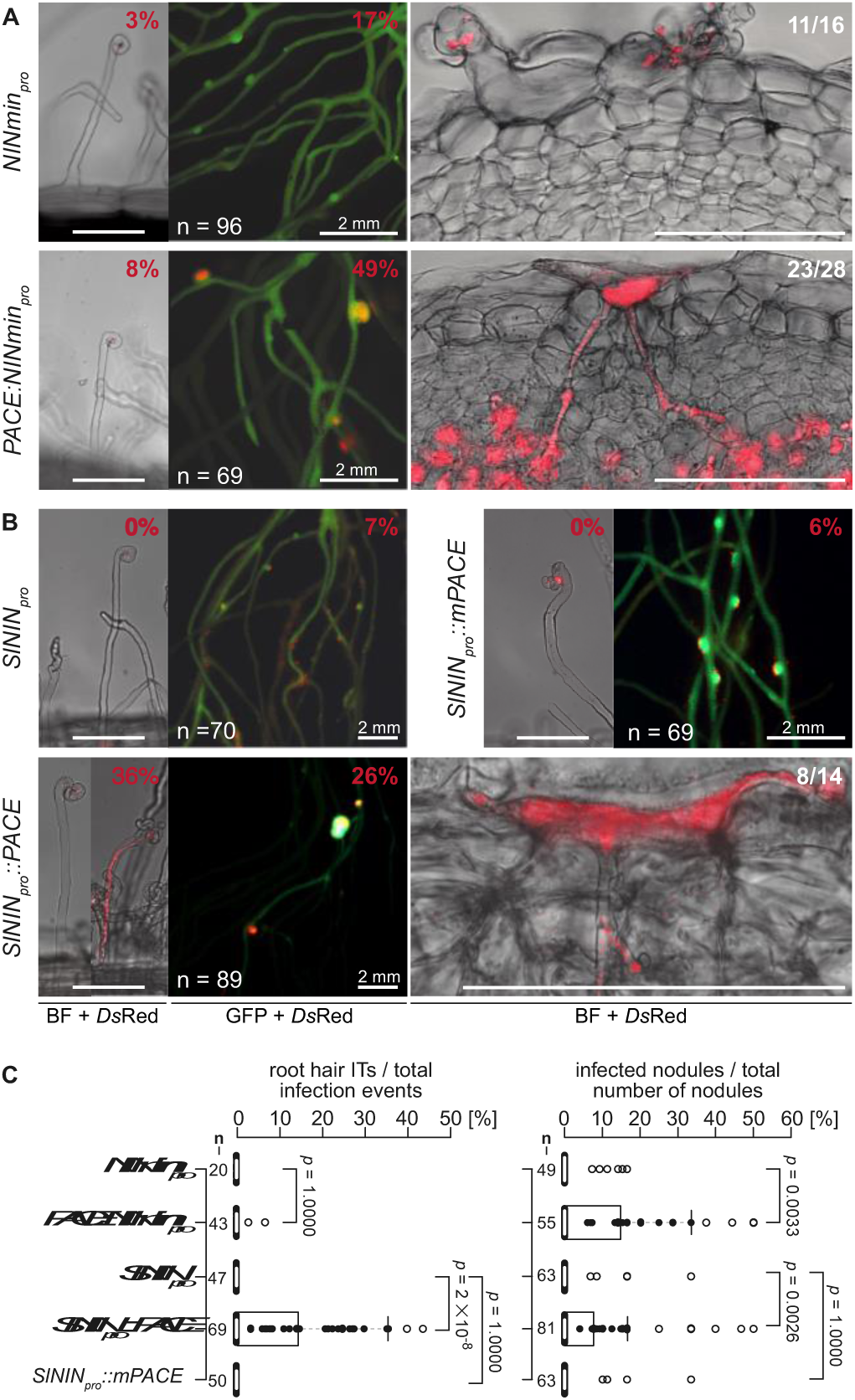
*PACE* enables IT formation in the cortex. Representative pictures of *nin-15* root hairs, root and nodule sections (see **Extended Data Fig. 10** for overview pictures), transformed with the *L. japonicus NIN* gene driven by (**A**) *NINmin_pro_* or *PACE:NINmin_pro_*; (**B**) *S. lycopersicum NIN* promoter (*SlNIN_pro_*) and *SlNIN_pro_* with *LjPACE* or *mPACE* inserted. %: percentage of transgenic root systems carrying root hair ITs or infected nodules; Ratios: number of nodules showing the presented pattern / total number of nodules sectioned and inspected. Data in (**A**) are from two independent experiments (see **Extended Data Fig. 10** for second replicate) and data in (**B**) are from a single experiment. (**C**) Boxplots displaying the percentage of root hair ITs and infected nodules per transgenic root system. Thick white lines, median; box, interquartile range; whiskers, lowest and highest data point within 1.5 interquartile range (IQR); black filled circles, data points inside 1.5 IQR; white filled circles, data points outside 1.5 IQR of the upper quartile. n, number of transgenic root systems or root pieces analysed. The applied statistical method was two-tailed Fisher’s exact test. Data in (**C**) are from (**A**) and (**B**). BF, brightfield. Unlabelled scale bars, 100 µm.

Transformation with the *L. japonicus NIN* gene driven by the *NIN* minimal promoter (*NINmin_pro_:NIN*) did not alter the symbiotic phenotype of *nin-15* roots (**Fig. 3B**). In contrast, the *NIN* gene driven by the *NIN* promoter (*NIN_pro_:NIN*) led to restoration of the complete infection process in *nin-15* roots from root hair ITs to symbiosome formation (100% and 92% of transgenic root systems carried root hairs ITs and infected nodules, respectively; **Fig. 3B**). Similar to observations in complementation experiments of *nin-2*, mutation or deletion of *PACE* (*NIN_pro_::mPACE:NIN* and *NIN_pro_:: ΔPACE:NIN,* respectively) drastically reduced the restoration of bacterial infection in root hairs and nodules in *nin-15* (**Fig. 3B**; **Extended Data Fig. 6 – 9** and **Supplementary Fig. 6 – 7**).

### *PACEs* from different nodulating FaFaCuRo species are functionally equivalent

*PACE* was detected by MEME searches as a conserved motif within *NIN* promoters of the FaFaCuRo clade. However, the individual *PACE* sequences from different species differed from each other, mostly so in the sequences flanking the *CYC-box* (**Fig. 1** and **Extended Data Fig. 1**). We therefore tested whether and to what extend this sequence variation of *PACE* would affect its function. Replacement of *PACE* within the *L. japonicus* (Fabales) 3 kb *NIN* promoter with *PACE* sequence variants (*NIN_pro_::Species abbreviation PACE:NIN*) originating from *Casuarina glauca* (*Cg*, Fagales), *Datisca glomerata* (*Dg*, Cucurbitales) or *Dryas drummondii* (*Dd*, Rosales) restored the complete infection process in *nin-15* to similar level as *NIN_pro_:NIN*, demonstrating the functional conservation of *PACE* from nodulating species across the entire FaFaCuRo clade (**Fig. 3C** and **Extended Data Fig. 9**). Similarly, the *PACE* versions from two non-nodulating Rosales that maintained the *NIN* gene, *Ziziphus jujuba* and *Prunus persica*, restored the complete infection process in *nin-15* (**Fig. 3D** and **Extended Data Fig. 9**). The results of these complementation experiments were consistent with the conserved expression pattern mediated by *PACEs* in *L. japonicus* (**Supplementary Fig. 8**) and the CCaMK/Cyclops-mediated transactivation via these *PACE* variants (**Extended Data Fig. 3A**) or chimeric promoter:reporter fusions (**Extended Data Fig. 3B**) tested in *N. benthamiana* leaves.

### Loss of *PACE* is associated with a loss of the nitrogen-fixing root nodule symbiosis

Griesmann et al. ^15^ and van Velzen et al. ^16^ discovered that RNS was lost multiple times independently during evolution, via independent truncations or losses of the *NIN* gene. However, at least 10 out of 28 FaFaCuRo species that lost RNS have maintained a full-length *NIN* open reading frame (**Supplementary Table 3**). Based on our complementation data, *PACE* is indispensable for the *NIN* promoter function in symbiosis (**Fig. 3**; **Extended Data Fig. 6 – 9** and **Supplementary Fig. 6 – 7**). Therefore, the absence of *PACE* from 5 out of these 10 species (**Supplementary Tables 1 – 3**), is potentially sufficient to explain these losses of RNS. Consequently, at least 82% of all losses can now be attributed to either the *NIN* ORF (18/28, 64%) or loss of *PACE* (5/28, 18%) (**Supplementary Table 3**). The presence of *PACE* in all nodulating species (31/31, 100%; **Supplementary Tables 1 and 2**) together with a correlation between the absence of *PACE* with the absence of RNS adds strong support for the evolutionary relevance of *PACE* both in the gain and potential loss of RNS.

*PACE* was not detected in the promoters of *NIN-like protein* (*NLP*) genes (**Supplementary Fig. 1** and **Supplementary Tables 1 and 2**) with the possible exception of the curious case of *Juglans regia* (Fagales). While it was also absent from the promoter of the so-annotated *NIN* gene, a *PACE*-like motif was identified in the promoter of the closest gene family member, *NIN-like protein 1 JrNLP1b* (*JrPACE-like*; **Supplementary Table 1**). This *PACE*-like element was not able to restore IT formation in *nin-15* (**Fig. 3D** and **Extended Data Fig. 9**). Regardless of whether this exceptional presence/absence pattern of *PACE* may be caused by a miss-annotation of *NIN* and *NLP1* in *J. regia*, either a loss-of-function mutation within *PACE* or a loss of the entire *PACE* element in the *JrNIN* promoter could explain the absence of the RNS observed in this species.

### *PACE* is sufficient to restore cortical IT formation in *nin-15*

We tested whether *PACE* on its own, only supported by the minimal *NIN* promoter (*PACE:NINmin_pro_*) is sufficient to restore IT development in cortical cells. For this purpose, we transformed *nin-15* roots with *PACE:NINmin_pro_* fused to the transcribed region of the *NIN* gene. *PACE*-mediated *NIN* expression led to an increased success in restoration of infection (49% of transgenic root systems carried infected nodules) compared to *NINmin_pro_:NIN*-transformed roots (17%; **Fig. 4A**; **Extended Data Fig. 10** and **Supplementary Table 4**). Root hair ITs were rarely observed on *PACE:NINmin_pro_:NIN*-transformed *nin-15* roots (**Fig. 4A**) and nodules harbouring cells filled with symbiosomes were not found (**Extended Data Fig. 10**), consistent with the restricted expression domain defined by *PACE* (**Extended Data Fig. 4**).

Strikingly, the vast majority of infected nodules transformed with *PACE:NINmin_pro_:NIN* (25 out of 28 nodules inspected) carried ITs in the outer cortex, originating from a focused hyperaccumulation of bacteria, locally constricted by root cell wall boundaries (**Fig. 4A**). This phenomenon was not observed in most of the rarely occurring infected nodules formed on *NINmin_pro_:NIN*-transformed *nin-15* roots (11 out of the 16 nodules inspected did not carry ITs in the outer cortex). Bacterial colonies within cell wall boundaries resembling this phenomenon have been described in a variety of legumes including *Sesbania* and *Mimosa* ^51,52^. Our data imply that *PACE* promotes this type of cortical IT initiation. Altogether, these findings revealed that *PACE* promotes IT development in cortex cells but not within root hairs.

### *PACE* insertion into the tomato *NIN* promoter confers RNS capability

To artificially recapitulate the functional consequence of *PACE* acquisition into a non-FaFaCuRo *NIN* promoter, we chose tomato (*Solanum lycopersicum*) which belongs to the Solanaceae, a family phylogenetically distant from the FaFaCuRo clade. Consistent with the absence of *PACE*, a *GUS* reporter gene driven by the tomato *NIN* promoter (*SlNIN_pro_*) was not transactivated by Cyclops in *N. benthamiana* leaf cells (**Fig. 1** and **Extended Data Fig. 2B and 3C**), while the insertion of the *L. japonicus PACE* (*SlNIN_pro_::PACE*), but not of a mutated *PACE* (*SlNIN_pro_::mPACE*) conferred transactivation by Cyclops (**Extended Data Fig. 3C**).

We tested the ability of the *LjNIN* expressed under the control of these synthetic promoters to restore the bacterial infection process in *nin-15*. Similar to *NINmin_pro_:NIN-*transformed *nin-15* roots, *SlNIN_pro_:NIN* did not restore bacterial infection (0% and 7% of transgenic root systems carried root hair ITs and infected nodules, respectively; **Fig. 4A – C**). In contrast, *nin-15* roots transformed with *SlNIN_pro_::PACE:NIN* restored the formation of root hair ITs and infected nodules on 36% and 26% of transgenic root systems, respectively (**Fig. 4B** and **Extended Data Fig. 10**). This increase in infection success was not observed on *SlNIN_pro_::mPACE:NIN*-transformed roots. ITs in the outer cortex that originated from a focal accumulation of bacteria were also observed in the *SlNIN_pro_::PACE:NIN*-transformed *nin-15* nodules (8 out of 14 nodules inspected; **Fig. 4B**) resembling those in the *PACE:NINmin_pro_:NIN*-transformed *nin-15* nodules (**Fig. 4A**). The gained ability of the *SlNIN::PACE* promoter to restore root hair ITs suggested that additional *cis*-regulatory elements within the *SlNIN* promoter function together with *PACE* for root hair IT formation. All together, these findings obtained with the tomato *NIN* promoter carrying an artificially inserted *PACE* agree with the hypothesis that the acquisition of *PACE* by a non-FaFaCuRo *NIN* promoter enabled its regulation via Cyclops and laid the foundation for IT formation in cortical cells.

## Discussion

The mechanistic connection between *PACE* and cortical IT formation together with their congruent phylogenetic distribution strongly support the idea that the acquisition of *PACE* by the latest common ancestor of the FaFaCuRo clade enabled cortical ITs and thus laid the foundation for the evolution of present day RNS. Our findings support an evolutionary model in which an ancestral symbiotic transcription factor complex (comprising CCaMK and Cyclops), that facilitated intracellular symbiosis with AM fungi already in the earliest land plants ^53,54^, gained control over the transcriptional regulation of the *NIN* gene by the acquisition of *PACE* (**Fig. 1**). This genetic innovation in the last common ancestor of the FaFaCuRo clade extended the function of the ancestral CCaMK complex to initiate cortical IT development. The *NIN*-like protein family underwent important evolutionary steps preceding the origin of RNS including a gene duplication leading to *NIN* and *NLP1* as closest paralogs ^55^. It is very likely that the NIN protein itself underwent changes that enabled its role in nodulation ^35^. Loss of *NIN* events associated with the loss of nodulation are scattered across all four FaFaCuRo orders ^15,16^, suggesting that *NIN* acquired its relevance for nodulation probably before or latest in the last common ancestor. Because our phylogenomic analysis dates the acquisition of *PACE* to the latest ancestor of the FaFaCuRo clade, we conclude that the critical changes within NIN must have occurred simultaneously or earlier. From a statistical point of view, it is likely that the *PACE* acquisition and the RNS-enabling changes within NIN occurred independently from each other. It will be interesting to determine what these critical changes within NIN are and where they occurred phylogenetically.

A “young” primary cell wall characteristic for recently divided cells is considered an important prerequisite for cortical IT initiation ^6,56^ but cell division is not restricted to the formation of novel organs ^9^. It is therefore conceptually possible that the common ancestor of the FaFaCuRo clade was forming ITs in recently divided cortical cells but in the absence of root nodules. Multiple lines of evidence indicate that the diverse types of lateral organs harbouring nitrogen-fixing bacteria (“nodules”) evolved multiple times independently. Indeed, *CE*-mediated *NIN* expression is important for nodule organogenesis in legumes, but upon searching for this regulatory element in a region of 0.1 Mb upstream and downstream of the *NIN* gene, Liu and colleagues ^40^ found its presence to be restricted to legume species, indicating an evolutionary emergence independently of and significantly later than the last common ancestor ^40,55^. ITs in root hairs are only found in Fabales and Fagales and therefore also considered a more recent acquisition ^6,57^. *CE* only in combination with *PACE* facilitates root hair ITs (**Extended Data Fig. 6 – 8** and **Supplementary Fig. 6 – 7**) and additional elements in the 3 kb promoter are necessary for nodule and cortical IT development (**Extended Data Fig. 6 – 8** and **Supplementary Fig. 6 – 7**). Furthermore, deletion of *PACE* by targeted genome editing was recently reported to reduce but not completely abolish the formation of root hair ITs suggesting the presence of partly redundant *PACE* elements in the vicinity of the *LjNIN* gene ^58^. Altogether, these observations highlight the complexity and concerted activity of *cis*-elements and transcription factors underlying the spatiotemporal expression control by present day *NIN* promoters in RNS-competent species. Our data pinpoint the acquisition of *PACE* as a key event during the evolution of the nitrogen-fixing root nodule symbiosis. Together with our discovery that multiple independent losses of *PACE* are associated with multiple losses of RNS within the FaFaCuRo clade, our data underpin the essential position of *PACE* in the evolutionary gain and loss of RNS.

## Materials and Methods

### Bioinformatic analyses

Based on the phylogenetic classification of the RWP-RK gene family ^15,50^, 144 NIN/NLP genes were selected from 37 plant species and 13 orders ranging from monocotyledons to dicotyledons including the FaFaCuRo clade (**Supplementary Table 1**). For each selected gene, 3 kb of sequence upstream of the translational start site including promoter and 5’UTR region was defined and extracted from the corresponding species’ genomic sequence, if contig length allowed it. For *Medicago truncatula,* a 3352 bp sequence upstream of the translational start site was extracted. If contig length was limiting, the longest possible sequence stretch was extracted. For the identification of a *cis*-regulatory element specific for *NIN* promoters of the FaFaCuRo clade, the tool MEME (http://meme-suite.org/tools/meme) was used in discriminative mode (option: “search given strand only”, default parameters) with *NIN* promoter regions of only nodulating plants. The control group consisted of promoter regions of all *NIN* genes outside of the FaFaCuRo clade and all NLP genes listed in **Supplementary Table 1**. The highest scoring motif (E-value: 1.6e-058) was 27 bp long and contained the much shorter previously described *CYC-box* ^34^ (**Extended Data Fig. 1A**).

To refine the conserved region in this motif, MEME analysis was performed again in normal mode (option: “search given strand only”, default parameters) with *NIN* promoters from only nodulating species. This analysis revealed that the most conserved nucleotides are found within 29 nucleotides (nucleotides 10 to 38 in **Extended Data Fig. 1B**). The previous MEME analysis was repeated, but an exactly 29 nucleotides long motif was searched for (resulting in a motif in **Extended Data Fig. 1C**) and the best scoring *NIN* paralog per searched species, i.e. lowest p-value per species were identified (**Supplementary Table 1**). In a final step, one best scoring *NIN* promoter region per nodulating species were analysed with MEME (option: “search given strand only”, default parameters) by searching for an exactly 29 nucleotides long motif. The resulting motif was named *PACE* (**Extended Data Fig. 1D**). This final MEME run was done for two reasons: first in order to avoid a sequence bias towards a single species with multiple *NIN* paralog promoter regions (e.g. soybean); second to avoid a potential sequence bias generated by the promoter region of a *NIN* paralog that might be no longer functional and therefore has mutated sites in its promoter region due to relaxed selection pressure.

As a control, the FIMO tool was used (http://meme-suite.org/tools/fimo, option: “scan given strand only”, default parameters, fdr < 0.1) to search all 144 *NIN* and *NLP* promoter regions (**Supplementary Table 1**). *PACE* was found within *NIN* promoter regions of all nodulating species analysed and two non-nodulating FaFaCuRo species (*Prunus persica* and *Ziziphus jujuba*) (**Supplementary Table 1**).

The presence or absence of *PACE* was further investigated in promoters of *NIN* and *NLPs* in an expanded database of 163 species encompassing 39 orders covering 6 groups of Viridiplantae (**Supplementary Table 2**). Orthologs of the whole *NLP* family were retrieved using tBLASTn v2.11.0+ ^59^ with reference sequences from *Medicago truncatula* as query and a cut-off e-value of 1e-10. Sequences were then aligned using MAFFT v7.380 ^60^ with default parameters. To identify the NIN and NLPs orthologues and therefore resolve the NIN and NLP protein subfamilies, we employed a Maximum Likelihood approach using the IQ-TREE v1.6.7 software ^61^. Prior to phylogenetic reconstruction, the best-fitting evolution model was determined for each alignment using ModelFinder ^62^ as implemented in IQ-TREE. Branch support was tested using 10,000 replicates of UltraFast Bootstraps using UFBoot2 ^63^. For each identified ortholog, a 5 kb region upstream of the translational start site was extracted. The three different consensuses identified in the previous MEME analyses (**Extended Data Fig. 1A, B, D**) were then searched in all *NLP* upstream regions using FIMO 5.0.2 and a q-value threshold of 0.1. If several motifs were identified in a given upstream region, only the one with the lowest q-value was conserved for further analysis (**Extended Data Fig. 1E**; **Supplementary Fig. 1** and **Supplementary Table 2**). *PACE* from *Parasponia andersonii* (a nodulating species from Rosales) was identified in this analysis and included to generate the consensus in **Fig. 1**.

Promoters of *ERN1* and *RAM1* genes were analysed independently of the previous analysis, using 87 plant genomes covering the main Angiosperms orders (**Supplementary Table 5**). Orthologs of each gene were retrieved using tBLASTn v2.7.1+ ^59^ with reference sequences from *Medicago truncatula* as query and a cut-off e-value of 1e-10. Sequences were then aligned using MAFFT v7.380 ^60^ with default parameters. Alignments were subjected to phylogenetic analysis to identify orthologs using Maximum Likelihood approach and the IQ-TREE v1.6.7 software ^61^. Prior to phylogenetic reconstruction, the best-fitting evolution model was determined for each alignment using ModelFinder ^62^ as implemented in IQ-TREE. Branch support was tested using 10,000 replicates of UltraFast Bootstraps using UFBoot2 ^63^. For each identified ortholog, regions upstream of the translational start site of different lengths (1 to 5 kb) were extracted. For each length of region upstream of the translational start site, sequences were analysed using MEME software v5.0.1 ^64^ with the following parameters: a motif size comprised between 5 - 45 bp and 5 - 25 bp for *ERN1* and *RAM1* respectively and a maximum number of discovered motifs of 20 (**Supplementary Fig. 2 and 3**). In addition, MEME search was set on “zoops” mode, assuming that each sequence can contain zero or one occurrence of the motif.

### Biological material

*L. japonicus* ecotype Gifu B-129 wild-type (WT) ^65^, *nin-2* ^32^ and *nin-15* (*LORE1* line 30003529 ^66^) were used in this study. Seed bags, bacterial strains and days post inoculation for each experiment are listed in **Supplementary Table 6**.

### Plant growth conditions and symbiotic inoculations

*Lotus japonicus* seeds were scarified and surface-sterilized as described ^67^ before germination on ½ Gamborg’s B5 medium solidified with 0.8 % Bacto™ agar in square plates (12 x 12 x 1.7 cm) ^68^. Plates were kept in dark for three days before transferring to light condition in a Panasonic growth cabinet (MLR-352H-PE) at 24 °C under a 16 h/8 h light/dark regime (50 µmol·m-2·s-1). Six-days-old seedlings were (1) subject to hairy root transformation as described ^69^ (**Fig. 2, 3 and 4** ; **Extended Data Fig. 4 – 10** and **Supplementary Fig. 5 – 8**); or (2) transferred to Weck jars (SKU 745 or 743; J.Weck GmbH u. Co. KG) containing 300 ml of sand:vermiculite mixture (2:1) and 20 ml of a modified ¼ strength Hoagland’s medium with Fe-EDDHA used as iron source ^70^ (**Extended Data Fig. 5E**). For *in vivo* promoter expression analysis (**Fig. 2** and **Supplementary Fig. 5**), transgenic roots expressing a kanamycin-resistance gene were kept on square plates supplemented with kanamycin (25 µg/ml) 10 days after *Agrobacterium rhizogenes* inoculation. Plants with transformed roots were kept on 0.8% Bacto™ agar including a nitrogen-reduced version of FAB medium (500 µM MgSO_4_·7H_2_O; 250 µM KH_2_PO_4_; 250 µM KCl; 250 µM CaCl_2_·2H_2_O; 100 µM KNO_3_; 25 µM Fe-EDDHA; 50 µM H_3_BO_3_; 25 µM MnSO_4_·H_2_O; 10 µM ZnSO_4_·7H_2_O; 0.5 µM Na_2_MoO_4_·2H_2_O; 0.2 µM CuSO_4_·5H_2_O; 0.2 µM CoCl_2_·6H_2_O; pH 5.7) in square plates for 1 week before transferring to a growth chamber at 24 °C under a 16 h/8 h light/dark regime (275 µmol·m-2·s-1) in Weck jars (SKU 745 or 743) containing 300 ml of sand:vermiculite mixture (2:1) and 30 ml of nitrogen-reduced FAB medium containing *Mesorhizobium loti* MAFF 303099 *Ds*Red ^71^ (*M. loti Ds*Red; **Fig. 3 – 4**; **Extended Data Fig. 5 – 10** and **Supplementary Fig. 6 – 7**), *M. loti* R7A CFP ^72^ (**Fig. 2**) or *M. loti* MAFF 303099 GFP ^73^ (**Supplementary Fig. 5**) set to a final optical density at 600 nm (OD_600_) of 0.05. For **Extended Data Fig. 4** and **Supplementary Fig. 8**, plants were grown in Weck jars (SKU 745 or 743) containing 300 ml of sand:vermiculite mixture (2:1) and 60 ml of nitrogen-reduced FAB medium containing *M. loti Ds*Red or MAFF 303099 *lacZ* ^72^ (*M. loti lacZ*) (OD_600_ = 0.01).

### Cloning and DNA constructs

For the construction of promoter:*NIN* fusions for complementation experiments (**Fig. 3**; **Extended Data Fig. 6 – 10** and **Supplementary Fig. 6 – 7**), the *NIN* genomic sequence without the 5’ and 3’UTRs served as a cloning module. A 3 kb region of the *L. japonicus NIN* promoter plus the 244 bp *NIN* 5’UTR was cloned from *L. japonicus* Gifu and used for complementation experiments (**Fig. 3**; **Extended Data Fig. 6 – 9** and **Supplementary Fig. 6 – 7**), dual luciferase assays (**Extended Data Fig. 2**), fluorimetric GUS assay (**Extended Data Fig. 3**) and promoter activity analysis (**Fig. 2**; **Extended Data Fig. 4** and **Supplementary Fig. 5**). For all the other versions of the *L. japonicus NIN* promoter tested (**Fig. 2, 3 and 4** ; **Extended Data Fig. 3 – 10** and **Supplementary Fig. 5 – 8**), the *LjNIN* minimal promoter (98 bp ^34^) plus the *LjNIN* 5’UTR was fused to 3’ end of the promoter. A 472 bp region containing multiple cytokinin response elements and highly conserved in eight legume species was identified 5’ of the *NIN* transcriptional start site by Liu et al. ^40^. We used this conserved region of 472 bp from *L. japonicus* and added flanking regions (192 bp upstream and 366 bp downstream; 2399 bp upstream and 2231 bp downstream, respectively) to obtain cytokinin element-containing regions of 1 kb and 5 kb (*CE_1kb_* and *CE_5kb_*, respectively). The *Solanum lycopersicum* gene ID Solyc01g112190.2.1 was identified as the closest homologue of *LjNIN* gene based on phylogenetic analysis ^15^, and is referred to as *SlNIN*. A 3 kb region of the *SlNIN* promoter plus the 238 bp *SlNIN* 5’UTR was cloned from *S. lycopersicum* cv. “Moneymaker” and *PACE* or *mPACE* (**Extended Data Fig. 3A**) was inserted 184 bp upstream of the *SlNIN* 5’UTR and used for complementation experiments (**Fig. 4** and **Extended Data Fig. 10**), dual luciferase assays (**Extended Data Fig. 2**) and fluorimetric GUS assay (**Extended Data Fig. 3**). A detailed description of constructs can be found in **Supplementary Fig. 4** and **Supplementary Table 7**. A list of oligonucleotides can be found in **Supplementary Table 8**. Constructs were generated with the Golden Gate cloning system ^74^.

### Imaging

Microscope and scanner settings as well as parameters for image acquisition are listed in Supplementary Table 9.

### Phenotypic analysis and quantification of infection events

Infected and non-infected nodules were discriminated by the presence and absence of a *Ds*Red signal (representing *M. loti Ds*Red) detected or not detected inside of the nodules, respectively. Presence or absence of bacteria was later confirmed by examination of sections of representative nodules. Infection threads (ITs) and *M. loti* entrapments in root hairs were detected by their *Ds*Red fluorescence (for microscope settings see **Supplementary Table 9**).

For phenotypic analysis of *nin-15* (**Extended Data Fig. 5C, F**), quantification was performed 21 days after inoculation (dpi) with *M. loti Ds*Red as follows: (1) the total number of nodules (including infected and non-infected) was determined under white light illumination (WLI); (2) the number of infected nodules and root hair ITs were counted as described above. Shoot dry weight was measured after drying the shoot at 60 °C for 1 h (**Extended Data Fig. 5E**).

For complementation experiments of *nin-2* and *nin-15* (**Fig. 3 and 4** ; **Extended Data Fig. 6 – 10** and **Supplementary Fig. 6 – 7**), quantifications and sectioning were performed 21 or 35 dpi with *M. loti Ds*Red with the microscope settings listed in **Supplementary Table 9** in the following order: (1) transgenic roots were identified by GFP fluorescence-emanating nuclei with a GFP filter; (2) infected nodules were counted as described above; (3) the total number of nodules (including infected and non-infected ones) was then determined under WLI; (4) the number of non-infected nodules was calculated by subtracting the number of infected nodules from the total number of nodules. To quantify infection events in root hairs, the number of bacterial entrapment and ITs in root hairs were counted on a 0.5 cm root piece for each transgenic root system, excised from a region where bacterial accumulation was detected by *Ds*Red fluorescence. Sectioning was performed on non-infected and infected nodules and the presence/absence of ITs and symbiosomes in cortical cells was examined. Nodule primordia and nodules were embedded in 6% low-melting agarose and sliced into 40 - 50 µm thick sections using a vibrating-blade microtome (Leica VT1000 S).

### Transient expression in *Nicotiana benthamiana* leaves

*Agrobacterium tumefaciens* strain AGL1 carrying promoter:reporter fusions on T-DNA were infiltrated as described in ^75^ with the acetosyringone concentration in the infiltration buffer modified to 150 µM. *A. tumefaciens* strains AGL1 and GV3101 containing plasmids *35S_pro_:3xHA-Cyclops* ^34^ and *35S_pro_:CCaMK*^1–314^–*mOrange* ^76^, respectively, were co-infiltrated with the reporter constructs as indicated. An AGL1 strain carrying a K9 plasmid constitutively expressing red fluorescent protein was used as needed to equalize the density of the *A. tumefaciens* suspension infiltrated per leaf, together with an *A. tumefaciens* strain carrying a plasmid for the expression of the viral P19 silencing suppressor to reduce post-transcriptional gene silencing ^77^ (**Extended Data Fig. 2 – 3**). *N. benthamiana* leaf discs with a diameter of 0.5 cm were harvested 60 hours post infiltration and used for quantitative fluorometric GUS assay and dual luciferase assay.

### Dual luciferase assay

The dual luciferase assay (**Extended Data Fig. 2**) was based on the Dual-Luciferase® reporter assay system (Promega). *N. benthamiana* leaf discs were ground to a fine powder in liquid nitrogen in 2 ml Eppendorf Safe-Lock tube (1 leaf disc in each tube) and subsequently incubated 5 min at room temperature (RT) with 200 µl of the Passive lysis buffer (Promega catalog number E1910). The resulting crude leaf extract was centrifuged at 20,000 g for 2 min at RT. An aliquot of the supernatant was subjected to the dual luciferase assay according to manufacturer’s instruction for Promega Dual Luciferase Kit (catalog number E1910) and chemiluminescence was quantified with a fluorescence plate reader (TECAN Infinite® 200 PRO; TECAN Group Ltd.) in white 96 well plates (Greiner Bio-One International GmbH). For each reporter construct, the promoter of interest was fused to the *Firefly* luciferase gene and constitutively expressed *Renilla* luciferase from the same vector was used for normalization (**Supplementary Table 7**). The ratio of the two signals (*Firefly* luciferase signal to the *Renilla* luciferase) was calculated and normalized to the vector control. A total number of at least 4 biological and 2 technical replicates per indicated vector were analysed in 2 independently performed assays.

### Quantitative fluorometric GUS assay and analysis

Quantitative fluorometric GUS assays (**Extended Data Fig. 3**) were performed as described ^78^ adapted to 96 well format. A total number of 7 to 8 leaf discs per indicated vector combination were analysed in 2 assays independently performed in different weeks.

### Promoter activity analysis

For promoter activity analyses with the *GUS* reporter gene (**Extended Data Fig. 4A – F** and **Supplementary Fig. 8**), transgenic nodule primordia and nodules were excised 10 - 14 dpi or ≥ 21 dpi with *M. loti Ds*Red and stained for GUS activity using 5-bromo-4-chloro-3-indolyl-beta-D-glucuronic acid (X-Gluc; x-gluc.com) as catalytic substrate ^75^ for 3 h at 37 °C. To visualise the root hair ITs together with the promoter activity with the *GUS* reporter gene (**Extended Data Fig. 4G**), plants with transgenic root systems were inoculated with *M. loti lacZ*. Transgenic roots were first stained for GUS activity with X-Gluc for 3 h at 37 °C and then for *lacZ* expression with Magenta-Gal for 18 h at 28 °C (as described in ^75^, which were visualized in blue and purple colour after staining, respectively). For promoter activity analyses with fluorescent reporters (**Fig. 2** and **Supplementary Fig. 5**), transgenic root systems were harvested 7 dpi (**Supplementary Fig. 5**) or 10 - 14 dpi (**Fig. 2**). Roots with bacterial infection at stage 2 - 3 and nodule primordia with bacterial infection at stage 3 or 4 (for stage description see main text) were selected by locating the GFP (*M. loti* MAFF 303099 GFP) and CFP signal (*M. loti* R7A CFP), respectively, via rapid (around 10 seconds) Z-stack analysis with the confocal light scanning microscope (**Supplementary Table 9**). For cell wall staining with calcofluor white (**Supplementary Fig. 5**), roots were fixed with 4% formaldehyde dissolved in PIPES buffer (50 mM Piperazine-N,N’-bis(2-ethanesulphonic acid), pH 7) for 1h under vacuum, rinsed 3 times with PIPES buffer and incubated in 0.05% calcofluor white dissolved in H_2_O for 1h. Roots and sections of nodule primordia were imaged as described in **Supplementary Table 9**.

### Data visualization and statistical analysis

Statistical analyses and data visualization were performed with RStudio 1.1. 383 (RStudio Inc.). Boxplots were used to display data in **Fig. 4**; **Extended Data Fig. 2, 3, 5, 6, 7, 9,10** and **Supplementary Fig. 6** (thick black or white lines: median; box: interquartile range; whiskers: lowest and highest data point within 1.5 interquartile range (IQR); black filled circles, data points inside 1.5 IQR; white filled circles, data points outside 1.5 IQR of the upper/lower quartile). The R package “beeswarm” with the method “center” was used to plot the individual data points for the boxplots (http://CRAN.R-project.org/package=beeswarm). The R package “agricolae” was used to perform ANOVA statistical analysis with post hoc Tukey and statistical results were displayed in small letters where different letters indicated statistical significance (https://cran.rproject.org/web/packages/agricolae/index.html). Tests applied are stated in the figure legend.

## Data availability

Raw data corresponding to **Fig. 1**, **Extended Data Fig. 1** and **Supplementary Fig. 1 – 3** are available in **Supplementary Tables 1, 2, 3 and 5**. The remaining raw data are available upon request. Essential plasmids listed in **Supplementary Table 7** can be ordered from the European Plasmid Repository (https://www.plasmids.eu/). References for the *L. japonicus* lines and *M. loti* strains are indicated in **Supplementary Table 6**.

## Acknowledgments

The authors thank David Chiasson for providing the MAFF 303099 *lacZ* and R7A CFP *M. loti* strains, Niels Sandal and Jens Stougaard for providing the *nin-15* mutant. This project has received funding from the European Research Council (ERC) under the European Union’s Seventh Framework Programme (FP7/2007-2013) under grant agreement n° 340904 (EvolvingNodules) which supported the work of MG and XG. MP acknowledges funding from the Deutsche Forschungsgemeinschaft (DFG) - Project number 170483403, in the context of the SFB924 ‘Molecular mechanisms regulating yield and yield stability in plants’ which supported the work of REA and CC and the ANR-DFG project ‘COME-IN’ Deutsche Forschungsgemeinschaft (DFG) – Project number 258665719, which supported the work of CC. This project has received funding from the European Union’s Horizon 2020 research and innovation programme under the Marie Sklodowska-Curie grant agreement No H2020-MSCA-IF-2015-703186 to KV and postdoctoral fellowship from Alexander von Humboldt Foundation to KV. PMD and JK belong to the LRSV laboratory, which is part of the TULIP Laboratoire d’Excellence (LABEX) (ANR-10-LABX-41). MH received funding from the JSPS KAKENHI Grant Number 17H06472.

## Author contributions

MG performed bioinformatic analysis of *NIN* promoters and discovered *PACE* presented in **Fig. 1** and **Extended Data Fig. 1A – D**. CC performed *in vivo* expression analysis presented in **Fig. 2** and **Supplementary Fig. 5** and prepared all confocal and light microscopy images of root hairs and nodule sections (**Fig. 3 and 4** ; **Extended Data Figu. 8 and 10** and **Supplementary Fig. 7**). Complementation experiments of *nin-15* were performed by REA (**Fig. 3B – D**, **Fig. 4A, 4C**; **Extended Data Fig. 9 and 10A – B)**, CC (**Fig. 3 and 4** ; **Extended Data Fig. 9C and 10) and XG (Fig. 3D and 4B – C**; **Extended Data Fig. 9 and 10C – D**). Complementation experiments of *nin-2* were performed by CC and XG (**Fig. 3A**; **Extended Data Fig. 6 – 8** and **Supplementary Fig. 6 – 7**). REA, CC and XG performed *nin-15* mutant phenotyping (**Extended Data Fig. 5**). XG drafted Supplementary Fig. 4 and performed transient expression assays presented in **Extended Data Fig. 3** and promoter expression analysis in **Extended Data Figure 4** and **Supplementary Fig. 8**. KV performed transient expression assays presented in **Extended Data Fig. 2** and drafted **Fig. 1**. JK and PMD performed motif search in *ERN1*, *NIN* and *RAM1* promoters presented in **Extended Data Fig. 1E** and **Supplementary Fig. 1 – 3**. CC, XG, REA, KV, PMD, MH, MG and MP formulated research hypothesis. CC, XG, REA, KV, MG and MP designed experiments. MP conceived and supervised the project. XG and MP coordinated research activities. MP, CC and XG wrote the manuscript and XG finalized all figures with inputs and comments from co-authors.

## Extended Data Figures

**Extended Data Figure 1 | Discovery of *PACE* by MEME analyses.** (**A - D**) Consensus sequence of the Position Weight Matrix identified by MEME analyses using the regions upstream of the translational start site (ATG) thus representing the promoters and 5’UTRs of *NIN* and *NLP* genes from 37 angiosperm species (see Methods): (**A**) in a discriminative search for a motif that is present in 3 kb upstream regions of the *NIN* genes from nodulating FaFaCuRo species, but absent in 3 kb upstream regions of the *NIN* genes from species outside of the FaFaCuRo clade and absent in the 3 kb upstream regions of *NLP* genes; (**B**) in an independent, non-discriminative search in the 3 kb upstream regions of the *NIN* genes from nodulating species revealing that the most conserved nucleotides span a larger region than (**A**); (**C**) the resulted most conserved 29 nucleotides derived from upstream regions of *NIN* genes of nodulating FaFaCuRo species; (**D**) the resulted most conserved 29 nucleotides (*PACE*) derived from the upstream region of one representative *NIN* gene per species of nodulating FaFaCuRo species. The Cyclops binding cite (*CYC-box* ^34^) is highlighted in grey. (E) Motif analyses by FIMO using the upstream regions of the *NIN* and *NLP* genes from an expanded list of 163 species (see Methods). Left: pruned tree from the whole *NLP* tree (demarcated in the black rectangle in **Supplementary Fig. 1**) corresponding to the *NIN* orthologs. Right: three versions of consensus sequence (left to right, (**A, B and D**), respectively) were used to retrieve motifs from the 5 kb upstream region of *NIN* genes *via* FIMO search, the output of which is displayed underneath the consensus sequences. Note that these identified motifs can originate from the + or – strand. The sequences displayed are those with the lowest q-value identified by FIMO and this resulted in some cases in overlapping sequences that originated from opposite strands (see **Supplementary Table 2**). Note that certain nodulating species have multiple copies of the NIN gene in their genome, for example *Phaseolus vulgaris* and *Glycine max*. We identified *PACE* in at least one of the *NIN* promoters in each of these species. Sequence names of nodulating species are coloured in blue. Blank lines represent the absence of significant motifs.

**Extended Data Figure 2 | Transcriptional activation of *NIN promoter:Firefly luciferase* reporter gene by CCaMK**^1–314^**/Cyclops is restricted to *NIN* promoters from species of the FaFaCuRo clade.** *Nicotiana benthamiana* leaf cells were transformed with T-DNAs carrying a *Firefly luciferase* reporter gene driven by either of the indicated promoters in tandem with the *AtACT2_pro_*:*Renilla luciferase* reporter fusion that provides a quantitative internal standard. (**A**) List of species within the FaFaCuRo clade (light red shade) and outside (light grey shade) and abbreviations. (**B**) Reporter gene activation by *L. japonicus* CCaMK^1–314^/Cyclops via *NIN* promoters (*NIN_pro_*) originating from listed species. (**C**) Comparison of the transactivation potential of Cyclops versions from *L. japonicus* and *S. lycopersicum*. Note that the expression of the *Firefly luciferase* reporter gene driven by *LjNIN_pro_*, the *RAM1* promoters from *L. japonicus* and *S. lycopersicum* (*LjRAM1_pro_*and *SlRAM1_pro_*, respectively) was induced in the presence of CCaMK^1–314^/Cyclops regardless of the origin of Cyclops. In contrast, the transactivation failed with the *SlNIN* promoter (panel (**A**)). Boxplots display the ratio of the Firefly/Renilla luciferase signals. Each dot represents one *N. benthamiana* leaf disc. Thick black lines, median; box, interquartile range; whiskers, lowest and highest data point within 1.5 interquartile range (IQR); black filled circles, data points inside 1.5 IQR; white filled circles, data points outside 1.5 IQR of the upper/lower quartile. The applied statistical method was ANOVA with *post hoc* Tukey: (**B**), *F_14,214_* = 71.07, *p* < 2×10^-16^; (**C**), plots from left to right: *F_5,18_* = 20.58, *p* = 7.14×10^-7^; *F_5,18_* = 25.38, *p* = 1.45×10^-7^ and *F_5,18_* = 40.49, *p* = 3.55×10^-9^, respectively. Different small letters indicate significant differences. Data displayed are from one experiment. Each combination of constructs was tested two times independently with similar outcomes.

**Extended Data Figure 3 | *PACE* sequence variants from species across the FaFaCuRo clade were able to functionally replace *L. japonicus PACE* in a *LjNIN_pro_:GUS* reporter fusion.** *N. benthamiana* leaf cells were transformed with T-DNAs carrying a *GUS* reporter gene driven by either of the indicated promoters: (**A**) the *L. japonicus NIN* promoter (*NIN_pro_*), the *LjNIN* promoter with *PACE* mutated or deleted (*NIN_pro_::mPACE* and *NIN_pro_::ΔPACE*, respectively), or *PACE* sequence variants from the nodulating FaFaCuRo species fused to the *LjNIN* minimal promoter (*NINmin_pro_*); (**B**) chimeric promoters where *LjPACE* in the *LjNIN* promoter was replaced with either one of the *PACE* variants from species tested in (**A**) or from non-nodulating FaFaCuRo species including the *Juglans regia PACE-*like motif (*JrPACE-like*); (**C**) the *S. lycopersicum NIN* promoter (*SlNIN_pro_*), the *SlNIN* promoter with *LjPACE* (*SlNIN_pro_::PACE*) or *mPACE* (*SlNIN_pro_::mPACE*) inserted. For species abbreviations see **Extended Data Fig. 2A**. Note in (**A**) that the deletion or mutation of *PACE* in *LjNIN* promoter resulted in a drastic reduction in reporter gene expression and in (**C**) insertion of *LjPACE* but not *mPACE* into the *S. lycopersicum* promoter confers transactivation by CCaMK^1–314^/Cyclops. GUS activities are displayed as individual dots in box plots. Each dot represents one *N. benthamiana* leaf disc. Thick black lines, median; box, interquartile range; whiskers, lowest and highest data point within 1.5 interquartile range (IQR); black filled circles, data points inside 1.5 IQR; white filled circles, data points outside 1.5 IQR of the upper/lower quartile. The applied statistical method was ANOVA with *post hoc* Tukey: (**A**) *F_20,144_*= 51.38, *p* < 2×10^16^; (**B**), *F_18,166_* = 149.1, *p* < 2×10^16^; (**C**) *F_7,62_* = 30.5, *p* = 7.02×10^-7^. Different small letters indicate significant difference. n.d., not determined. Data displayed are from one experiment. Each combination of constructs was tested two times independently with similar outcomes.

**Extended Data Figure 4 | Spatio-temporal *GUS* expression driven by *PACE* and the *NIN* promoter in *L. japonicus* roots during the bacterial infection process.** *L. japonicus* wild-type hairy roots were transformed with T-DNAs carrying a *Ubq10_pro_:NLS-GFP* transformation marker together with a *GUS* reporter gene driven by either of the indicated promoters: (**A**) the 3 kb *LjNIN* promoter (*NIN_pro_*); the *LjNIN* promoter with *PACE* (**B**) mutated (*LjNIN_pro_::mPACE*) or (**C**) deleted (*NIN_pro_::ΔPACE*); (**D**) *PACE* fused to the *LjNIN* minimal promoter (*PACE:NINmin_pro_*) or (**E**) the *LjNIN* minimal promoter (*NINmin_pro_*). The progression of bacterial infection was determined by the *Ds*Red signal 10 - 14 days post inoculation (dpi) with *M. loti Ds*Red. Nodules undergoing different stages of infection (panels I to IV) were stained with X-Gluc to reveal the *GUS* expression pattern. Note the overlapping bacterial invasion zone and *PACE:NINmin_pro_:GUS* expression in early infection stages (red and blue arrowheads in (**D**)) as well as the differences between *PACE:NINmin_pro_:GUS* and the much broader *NIN_pro_:GUS* expression at that stage (red and blue arrows in (**A**)). Red arrow and arrowheads: *M. loti Ds*Red. Blue arrow and arrowheads: GUS activity in root hairs bearing ITs and nodule primordia, respectively. The *NINmin_pro_*:*GUS* fusion gave only rarely detectable signal, and if so in the vasculature (yellow arrowhead in (**E**)). Only pictures taken under white light illumination (WLI) are displayed for nodules in panel VI to reveal the pink colour of leghemoglobin, characteristic for mature and fully infected nodules. Note that *PACE:NINmin_pro_:GUS* expression was absent at this stage, whereas the *NIN_pro_:GUS* resulted in strong blue staining in the nodule regardless of the presence of *PACE* (compare panel IV in (**D**) and (**A - C**)). (**F**) Quantification of transgenic root systems exhibiting *GUS* expression in different cell types and tissues exemplarily displayed in (**A - E**). (**G**) *PACE* drove *GUS* reporter gene expression in the central tissue of primordia and nodules, but was not sufficient for expression in root hairs. Transgenic roots carrying promoter:GUS fusions same as in (**A, D and E**) were inoculated with *M. loti lacZ* and dual-stained with X-Gluc and Magenta-Gal. Purple: *M. loti lacZ*. Blue: GUS activity. Note the co-existence of blue and purple staining in root hairs on roots transformed by *NIN_pro_:GUS*, but not that transformed by *PACE:NINmin_pro_:GUS*. Data displayed in (**F**) are combined from three independent experiments. Bars, 250 μm.

**Extended Data Figure 5 | *L. japonicus nin-15* mutant phenotype.** (**A**) A representative picture of *L. japonicus* wild-type (WT, left) and *nin-15* (right) plants 21 dpi with *M. loti Ds*Red. (**B**) Position of the *Lotus Retrotransposon 1* (*LORE1*) insertion within the *NIN* promoter in the *nin-15* mutant. (**C**) Representative pictures of *nin-15* root hairs and nodule sections 21 dpi with *M. loti Ds*Red. Forty-nine plants with a total number of 436 nodules were analysed: only four plants bore one or two IT(s) within root hairs and seven plants bore one or two infected nodule(s). Deformed or curled root hairs in the presence of *M. loti Ds*Red were abundant but infection threads were rarely found. Arrowheads: uninfected nodules. Unlabelled bars, 100 μm. (**D – E**) Phenotype of *nin-15* in the presence of a symbiosis-independent nitrogen source (15 mM KNO_3_) for 28 days. (**D**) Pictures documenting the healthy status of *L. japonicus* WT and *nin-15* plants (compare (**D**) and (**A**)) and (**E**) quantitative assessment of parameters displayed in boxplots. Thick black lines, median; box, interquartile range; whiskers, lowest and highest data point within 1.5 interquartile range (IQR); black filled circles, data points inside 1.5 IQR; white filled circles, data points outside 1.5 IQR of the upper/lower quartile. Each dot represents one plant. n: number of plants analysed. Lateral root density: number of lateral roots/primary root length (cm). The applied statistical method was two-tailed Welch’s *t-*test. (F) Segregation analysis of *nin-15* assessed by quantifying the number of infected nodules. Each dot in the boxplots represents one plant. n: number of plants analysed. The applied statistical method was ANOVA with *post hoc* Tukey: *F_3,120_* = 84.1, *p* = 2×10^16^. Different small letters indicate significant difference. (**G**) Representative pictures of *nin-15* plants with hairy roots transformed with the *NIN* gene driven by the *L. japonicus NIN* minimal promoter (*NINmin_pro_*) or the 3 kb *NIN* promoter (*NIN_pro_*) 24 dpi with *M. loti Ds*Red. Data are from a single experiment. WLI: white light illumination.

**Extended Data Figure 6 | The *CYC-box* and flanking sequences of *PACE* are required for the complete restoration of the bacterial infection process in the *L. japonicus nin-2* mutant.** *nin-2* roots were transformed with T-DNAs carrying a *Ubq10_pro_:NLS-GFP* transformation marker in tandem with the *LjNIN* gene driven by either of the following promoter versions: the cytokinin element-containing region of 1 kb (*CE_1kb_*) fused to the 3 kb or 9 kb *LjNIN* promoter (*CE_1kb_:NIN_pro_*or *CE_1kb_:NIN_9kbpro_*, respectively); *CE_1kb_:NIN_pro_*or *CE_1kb_:NIN_9kbpro_* with *PACE* mutated (*CE_1kb_:NIN_pro_::mPACE* or *CE_1kb_:NIN_9kbpro_::mPACE*, respectively); *CE_1kb_:NIN_pro_*carrying a mutated Cyclops binding site (*CYC-box*) (*CE_1kb_:NIN_pro_::mbox*); *CE_1kb_:NIN_pro_* carrying mutated sequences flanking the *CYC-box* in *PACE* (*CE_1kb_:NIN_pro_::mflanking*); *CE_1kb_* fused to the *LjNIN* minimal promoter (*CE_1kb_:NINmin_pro_*); *CE_1kb_* fused to *PACE* and to *NINmin_pro_* (*CE_1kb_:PACE:NINmin_pro_*); *NIN_pro_*, *PACE:NINmin_pro_* or *NINmin_pro_*. (**A**) Representative overview pictures of transgenic root systems. Roots were analysed 21 dpi with *M. loti Ds*Red. White asterisks and arrowheads: infected and non-infected nodules, respectively. Bars, 2 mm. (**B – C**) Boxplots displaying the number of root hair ITs or infected nodules and the percentage of root hair ITs among total infection events (sum of bacterial entrapments and ITs). Each dot represents one transgenic *nin-2* root system or root piece. *L. japonicus* WT roots transformed with *NIN_pro_:NIN* or *CE_1kb_:NIN_pro_:NIN* were included as controls. Note the loss of restoration of nodules and IT formation associated with the mutation of *PACE* or only the *CYC-box* in *PACE*; and the reduction of same when sequences flanking the *CYC-box* in *PACE* were mutated. n: number of transgenic root systems or root pieces analysed. Thick black lines, median; box, interquartile range; whiskers, lowest and highest data point within 1.5 interquartile range (IQR); black filled circles, data points inside 1.5 IQR; white filled circles, data points outside 1.5 IQR of the upper/lower quartile. Numbers above the boxplots: the value of individual data points outside of the plotting area. Data are from a single experiment. n.d.: not determined. WLI: white light illumination.

**Extended Data Figure 7 | The *CYC-box* and flanking sequences of *PACE* are required for the complete restoration of the bacterial infection process but are dispensable for the nodule organogenesis process in the *L. japonicus nin-2* mutant.** *nin-2* roots were transformed with T-DNAs carrying a *Ubq10_pro_:NLS-GFP* transformation marker in tandem with the *LjNIN* gene driven by either of the following promoter versions: the cytokinin element-containing region of 5 kb (*CE_5kb_*) fused to the 3 kb *LjNIN* promoter (*CE_5kb_:NIN_pro_*); *CE_5kb_:NIN_pro_* with *PACE* mutated (*CE_5kb_:NIN_pro_::mPACE*); *CE_5kb_:NIN_pro_* carrying a mutated Cyclops binding site (*CYC-box*) (*CE_5kb_:NIN_pro_::mbox*); *CE_5kb_:NIN_pro_* carrying mutated sequences flanking the *CYC-box* in *PACE* (*CE_5kb_:NIN_pro_::mflanking*); *CE_5kb_* fused to the *LjNIN* minimal promoter (*CE_5kb_:NINmin_pro_*); *CE_5kb_* fused to *PACE* and to *NINmin_pro_* (*CE_5kb_:PACE:NINmin_pro_*) or *NIN_pro_*. (**A**) Representative overview pictures of transgenic root systems. Roots were analysed 21 dpi with *M. loti Ds*Red. White asterisks and arrowheads: infected and non-infected nodules, respectively. Bars, 2 mm. (**B**) Boxplots displaying the number of infected nodules, the percentage of infected nodules among total organogenesis events (sum of infected and non-infected nodules) and the number of organogenesis events. (**C**) Boxplots displaying the number of root hair ITs and the percentage of root hair ITs among total infection events (sum of bacterial entrapments and ITs). Each dot represents one transgenic *nin-2* root system or root piece. *L. japonicus* WT roots transformed with *NIN_pro_:NIN* or *CE_5kb_:NIN_pro_:NIN* were included as controls. Note that the mutation of *PACE* or only the *CYC-box* in *PACE* led to an almost complete loss of IT formation and infected nodules per root system while nodule organogenesis was not significantly reduced; and that mutation of sequences flanking the *CYC-box* in *PACE* led to a reduction of the number of infected nodules per root systems. n: number of transgenic root systems or root pieces analysed. Thick black lines, median; box, interquartile range; whiskers, lowest and highest data point within 1.5 interquartile range (IQR); black filled circles, data points inside 1.5 IQR; white filled circles, data points outside 1.5 IQR of the upper/lower quartile. Data are from a single experiment. n.a.: not applicable. WLI: white light illumination.

**Extended Data Figure 8 | The *CYC-box* and flanking sequences of *PACE* are required for the complete restoration of the bacterial infection process but are dispensable for the nodule organogenesis process in the *L. japonicus nin-2* mutant.** Pictures of nodule sections or roots from *L. japonicus nin-2* roots 21 dpi with *M. loti Ds*Red from the same experiments depicted in **Extended Data Fig. 6** (**A**) and **Extended Data Fig. 7** (**B**). Nodule sections from *L. japonicus* WT roots transformed with *NIN_pro_:NIN* and *CE_5kb_NIN_pro_:NIN* were included for comparison. Note that when the cytokinin element-containing region of 1 kb was fused to *NIN_pro_* nodule organogenesis was abolished by mutation of *PACE* or only the *CYC-box* in *PACE* and that these mutations did not abolish organogenesis when the cytokinin element-containing region of 5 kb was fused to *NIN_pro_*. At least five nodules from independent transgenic root systems were sectioned per construct. Bars, 100 μm.

**Extended Data Figure 9 | *PACEs* from FaFaCuRo species are functionally equivalent in restoring bacterial infection in the *L. japonicus nin-15* mutant.** *L. japonicus nin-15* roots were transformed with T-DNAs carrying a *Ubq10_pro_:NLS-GFP* transformation marker in tandem with the *LjNIN* gene driven by either of the following promoters: (**A**) the 3 kb *LjNIN* promoter (*NIN_pro_*), the *LjNIN* minimal promoter (*NINmin_pro_*), the 3 kb *LjNIN* promoter with *PACE* deleted (*NIN_pro_::ΔPACE*) or mutated (*NIN_pro_::mPACE*); (**B**) the 3 kb *LjNIN* promoter with *LjPACE* replaced with either of the *PACE* sequence variants from nodulating or non-nodulating FaFaCuRo species and analysed 21 dpi with *M. loti Ds*Red. (**A – B**) Representative overview pictures of *nin-15* transgenic roots systems. Sections of representative nodules are displayed in **Fig. 3**. Note the drastic reduction of restoration of infection in nodules and root hairs associated with the mutation or deletion of *PACE* as well as the replacement of *PACE* with *JrPACE-like* in the context of the *LjNIN* promoter. White asterisks and arrowheads: infected and non-infected nodules, respectively. (**C – E**) Boxplots displaying (**C**) the percentage of root hair ITs among total infection events (sum of bacterial entrapments and ITs) and (**D – E**) the number of infected nodules from two independent experiments. Each dot represents one *nin-15* transgenic root piece (**C**) or root system (**D – E**). (**C**) displays merged data from experiments in (**D – E**) as the percentage represents a normalised value calculated for each root piece (see **Supplementary Table 4**). Thick black lines, median; box, interquartile range; whiskers, lowest and highest data point within 1.5 interquartile range (IQR); black filled circles, data points inside 1.5 IQR; white filled circles, data points outside 1.5 IQR of the upper/lower quartile. n: number of transgenic root systems or root pieces analysed. For species abbreviations see **Extended Data Fig. 2A**. The applied statistical method was ANOVA with *post hoc* Tukey: (**C**) *F_9,313_* = 106.7, *p* < 2×10^-16^; (**D**) *F_6,346_* = 82.89, *p* < 2×10^-16^; (**E**) *F_4,135_* = 20.18, *p* = 4.76×10^-13^. Different small letters indicate significant differences. Data are from a single experiment. Bars, 2 mm. WLI: white light illumination.

**Extended Data Figure 10 | *PACE* alone or in the context of the *S. lycopersicum NIN* promoter (a species outside of the FaFaCuRo clade) enables IT formation in the cortex.** (**A – D**) Representative pictures of sections of nodules formed on *L. japonicus nin-15* roots transformed with T-DNAs carrying a *Ubq10_pro_:NLS-GFP* transformation marker together with the *LjNIN* gene driven by either of the following promoters: (**A – B**) the *L. japonicus NIN* minimal promoter (*NINmin_pro_*) or *PACE* fused to *NINmin_pro_* (*PACE:NINmin_pro_*); (**C – D**) the 3 kb *S. lycopersicum NIN* promoter (*SlNIN_pro_*), the 3 kb *SlNIN* promoter with mutated *PACE* (*SlNIN_pro_::mPACE*) or with *L. japonicus PACE* inserted (*SlNIN_pro_::PACE*), 21 dpi with *M. loti Ds*Red (from the same experiments depicted in **Fig. 4**). Black rectangles in (**A**) demarcate the enlarged area displayed in **Fig. 4A and 4B** to focus on the initial infection structures. Note the absence of cells filled with symbiosomes in nodules transformed with the *LjNIN* gene driven by *PACE:NINmin_pro_*or *NINmin_pro_*. By contrast, infected cells were often filled with symbiosomes in the *SlNIN_pro_::PACE:NIN-*transformed nodules, like those resulted by *NIN_pro_:NIN* (see (**C**) and compare the two sections in (**D**)). (**E – F**) Boxplots displaying the percentage of root hair ITs among total infection events (sum of bacterial entrapments and ITs) or the percentage of infected nodules among total number of nodules (**E**) 21 dpi and (**F**) 35 dpi with *M. loti Ds*Red, respectively. Each dot represents one *nin-15* transgenic root system or root piece. (**E**) displays results from an independent repetition from the experiment depicted in **Fig. 4**. n: number of transgenic root systems or root pieces analysed. Thick white (**E**) and black (**F**) lines, median; box, interquartile range; whiskers, lowest and highest data point within 1.5 interquartile range (IQR); black filled circles, data points inside 1.5 IQR; white filled circles, data points outside 1.5 IQR of the upper/lower quartile. Data displayed are from a single experiment. Numbers above the boxplots: the value of individual data points outside of the plotting area. The applied statistical method was two-tailed Fisher’s exact test. Bars, (**A and C**) 100 μm; (**B and D**) 50 μm.

## Supplementary Figures

**Supplementary Figure 1 | PACE is present exclusively in the FaFaCuRo clade.** Motif analyses by FIMO using the using the regions upstream of the translational start site (ATG) thus representing the promoters and 5’UTRs of the *NIN* and *NLP* genes from an expanded list of 163 species (see Methods). Left: Maximum likelihood tree of *NLP* family (model: GTR+F+R10) of 163 species (**Supplementary Table 2**). Tree was rooted on the Charophytes algae *Klebsormidium nitens*. Black box demarcates the area that is displayed in **Extended Data Fig. 1E**. Blank lines represent the absence of significant motifs. Sequence names of nodulating species are coloured in blue. The different NLP clades are indicated by branch colours and named according to the *Arabidopsis thaliana* nomenclature. Right: three versions of consensus sequence (left to right, **Extended Data Fig. 1A, 1B and 1D**, respectively) were used to retrieve motifs from the 5 kb upstream region of *NIN* genes via FIMO search, the output of which is displayed underneath the consensus sequences.

**Supplementary Figure 2 | The presence of a motif encompassing the previously identified Cyclops binding site within the regions upstream of the translational start site (ATG) thus representing the promoters and 5’UTRs of *ERN1* genes extends beyond the FaFaCuRo clade.** (**A**) Maximum likelihood of *ERN1/ERN2* (model: JTT+F+R5) from 87 species (**Supplementary Table 5**) based on protein sequences. Tree was rooted using sequences from the *ERN1/ERN2* paralog *ERN3* (outgroup clade collapse and indicated by grey triangle marked “Outgroup”). Species belonging to the monophyletic FaFaCuRo clade (Stevens, http://www.mobot.org/MOBOT/research/APweb/) are indicated by thicker black branches. *ERN1* and *ERN2* subclades are indicated. Sequence names of nodulating species are coloured in blue. These conserved motifs identified by MEME analyses in the 3 kb upstream regions were aligned and displayed next to species names. While the presence of a conserved Cyclops binding site containing motif in species outside of the FaFaCuRo clade is consistent with the role of *RAM1* in arbuscular mycorrhizal symbiosis, a function of *ERN1* in arbuscular mycorrhizal symbiosis has not been described and suggests a yet unidentified function of *ERN1* outside of the FaFaCuRo clade. (**B**) Consensus sequence of the Position Weight Matrix identified by MEME analyses of all the species. (**C**) Consensus sequence of the Position Weight Matrix identified by MEME analyses when analysing only the nodulating species.

**Supplementary Figure 3 | The presence of a motif encompassing the previously identified Cyclops binding site within the regions upstream of the translational start site (ATG) thus representing the promoters and 5’UTRs of *RAM1* genes extends beyond the FaFaCuRo clade.** (**A**) Maximum likelihood of *RAM1* (model: JTT+F+R4) from 87 species (**Supplementary Table 5**) engaging in arbuscular mycorrhizal symbiosis. Tree was rooted on the early-diverging angiosperm *Amborella trichopoda*. Species belonging to the monophyletic FaFaCuRo clade (Stevens, http://www.mobot.org/MOBOT/research/APweb/) are indicated by thicker black branches. Sequence names of nodulating species are coloured in blue. These conserved motifs identified by MEME analyses in the 3 kb upstream regions are aligned and displayed next to species names. Blank lines represent the absence of identified motif due to genome contiguity issues. (**B**) Consensus sequence of the Position Weight Matrix identified by MEME analyses when analysing all the mycorrhizal species. (**C**) Consensus sequence of the Position Weight Matrix identified by MEME analyses when analysing only the nodulating species.

**Supplementary Figure 4 | Schematic representation of the *NIN* promoter and T-DNA constructs used in this study.** (**A**) Schematic representation of the *L. japonicus NIN* promoter and *cis*-regulatory regions that control *NIN* expression and enable rhizobia infection and nodule development. The *CE_1kb_* and *CE_5kb_* regions encompass several putative cytokinin response elements ^40^. The 3kb long promoter region encompasses *PACE* as well as the NSP1 and IPN2 binding sites ^47,48^. Numbers indicate the number of bases from the transcriptional start site (TSS). (**B - E**) Schematic drawing of the T-DNA region of plasmids used in different experiments. Details of these plasmids are listed in **Supplementary Table 7**.

**Supplementary Figure 5 | *PACE* drives the expression of the mCherry fluorescent reporter in cortical cells during IT development**. Representative pictures of *L. japonicus* root upon inoculation with *M. loti* MAFF 303099 expressing GFP (green) imaged by confocal laser-scanning microscopy. Comparison of the expression domains determined by (**A**) *PACE* (*PACE:NINmin_pro_:NLS-mCherry*; red) and (**B**) a *NIN* promoter carrying a mutated *PACE* (*NIN_pro_::mPACE:NLS-YFP*; yellow). The plant cell wall was stained with calcofluor white (cyan). Dashed lines indicate the membrane of the root hair cell. Arrowheads indicate the IT. Numbers: roots showing the presented expression pattern / total number of roots inspected. Data are from a single experiment. Bars, 20 μm.

**Supplementary Figure 6 | The *CYC-box* and flanking sequences of *PACE* are required for the complete restoration of the bacterial infection process in the *L. japonicus nin-2* mutant.** Roots were from a subset of plants from the same experiment depicted in **Extended Data Fig. 6** but analysed 35 dpi with *M. loti Ds*Red. (**A – B**) Boxplots displaying the number of root hair ITs or infected nodules and the percentage of root hair ITs among total infection events (sum of bacterial entrapments and ITs). Each dot represents one transgenic *nin-2* root system or root piece. *L. japonicus* WT roots transformed with *NIN_pro_:NIN* or *CE_1kb_:NIN_pro_:NIN* were included as controls. Note that the results follow the same trend as those obtained 21 dpi with *M. loti Ds*Red (**Extended Data Fig. 6**). n: number of transgenic root systems or root pieces analysed. Thick black lines, median; box, interquartile range; whiskers, lowest and highest data point within 1.5 interquartile range (IQR); black filled circles, data points inside 1.5 IQR; white filled circles, data points outside 1.5 IQR of the upper/lower quartile. Numbers above the boxplots: the value of individual data points outside of the plotting area. Data are from a single experiment.

**Supplementary Figure 7 | The *CYC-box* and flanking sequences of *PACE* are required for the complete restoration of the bacterial infection process in the *L. japonicus nin-2* mutant.** Pictures of nodule sections or roots from *L. japonicus nin-2* roots 35 dpi with *M. loti Ds*Red from the same experiments depicted in **Supplementary Fig. 6**. Upper left corner: a nodule section from a *L. japonicus* WT root transformed with *NIN_pro_:NIN* was included for comparison. At least five nodules from independent transgenic root systems were sectioned per construct. Bars, 100 μm.

**Supplementary Figure 8 | Spatio-temporal *GUS* expression driven by *PACE* variants in *L. japonicus* roots during the bacterial infection process.** *L*. *japonicus* WT roots were transformed with T-DNAs carrying a *Ubq10_pro_:NLS-GFP* transformation marker together with a *GUS* reporter gene driven by either of the *PACE* variants from nodulating FaFaCuRo species fused to the *LjNIN* minimal promoter (*NINmin_pro_*). For species abbreviations and experimental details see **Extended Data Fig. 2A and 4**, respectively. Note the overlapping bacterial invasion zone and *PACE:NINmin_pro_:GUS* expression in early infection stages (red and blue arrowheads in (**A – C**)). Red arrowheads: *M. loti Ds*Red. Blue arrowheads: GUS activity in nodule primordia. Only pictures taken under white light illumination (WLI) are displayed for nodules in panel VI to reveal the pink colour of leghemoglobin, characteristic for mature and fully infected nodules. Note that like *LjPACE*, the *PACE* variants-driven *GUS* expressions were absent at this stage (panel IV in (**A – C**) and panel IV in **Extended Data Fig. 4D**). (**D**) Quantification of transgenic root systems exhibiting *GUS* expression in different cell types and tissues exemplarily displayed in (**A – C**). n.d.: not determined. Data displayed in (**D**) are combined from two independent experiments. Bars, 250 μm.

## Supplementary Tables

**Supplementary Table 1 |** Summary of the bioinformatic analysis resulting in the discovery of *PACE* using 37 species.

**Supplementary Table 2 |** Results of the FIMO analysis of *PACE* in 163 species.

**Supplementary Table 3 |** Status of *PACE* and *NIN* in non-nodulating FaFaCuRo species.

**Supplementary Table 4 |** Results of hairy root mediated complementation experiments of the *L. japonicus* Gifu *nin-2* and *nin-15* mutant lines with indicated constructs.

**Supplementary Table 5 |** List of plant genomes used for the search of conserved motifs within *ERN1* and *RAM1* promoters.

**Supplementary Table 6 |** List of seed bags, bacterial strains and incubation times.

**Supplementary Table 7 |** List of plasmids used.

**Supplementary Table 8 |** Sequences and IDs of oligonucleotides (DNA) used.

**Supplementary Table 9 |** Microscope/scanner settings and image analysis.

